# Integrative analysis of mRNA stability regulation uncovers a metastasis-suppressive program in breast cancer

**DOI:** 10.1101/2025.06.06.658309

**Authors:** Heather Karner, Tabea Mittmann, Vicky W. Chen, Ashir A. Borah, Andreas Langen, Hassan Yousefi, Lisa Fish, Balyn W. Zaro, Albertas Navickas, Hani Goodarzi

**Affiliations:** Department of Biochemistry & Biophysics, University of California San Francisco, San Francisco, CA, USA; Department of Urology, University of California San Francisco, San Francisco, CA, USA; Helen Diller Family Comprehensive Cancer Center, University of California San Francisco, San Francisco, CA, USA; Faculty of Medicine, University of Münster, Münster, Germany; Biological and Medical Informatics Graduate Program, University of California San Francisco, San Francisco, CA, USA; Arc Institute, Palo Alto, CA, USA; Department of Genetics, Stanford University, Palo Alto, CA, USA; Department of Pharmaceutical Chemistry, University of California San Francisco, San Francisco CA, USA; Cardiovascular Research Institute, University of California San Francisco, San Francisco, CA, USA

## Abstract

Heterogeneity in cancer gene expression is typically linked to genetic and epigenetic alterations, yet post-transcriptional regulation likely influences these patterns as well. However, the quantitative contribution of post-transcriptional mechanisms to cancer transcriptome dynamics remains unclear. Here, we systematically measured mRNA dynamics across diverse breast cancer models, revealing that mRNA stability significantly shapes gene expression variability. To decipher the regulatory grammar underlying these dynamics, we developed GreyHound, an interpretable multimodal deep-learning framework integrating RNA sequence features and RNA-binding protein (RBP) expression. GreyHound identified an extensive network of RBPs and their regulons underlying variations in mRNA stability. Among these, we uncovered a metastasis-suppressive regulatory axis centered on the RNA-binding protein RBMS3 and its post-transcriptional control of the redox regulator TXNIP. Functional and molecular analyses revealed that RBMS3 depletion resulted in targeted transcript destabilization, which was associated with poor clinical outcomes and enhanced metastatic potential in xenograft models. Using in vivo epistasis studies, we confirmed that RBMS3 regulation of TXNIP mRNA stability drives this metastasis-suppressive program. These findings position the RBMS3-TXNIP regulatory axis as a key post-transcriptional mechanism in breast cancer and illustrate how interpretable models of RNA dynamics can uncover hidden regulatory programs in disease.

## Introduction

Precise regulation of gene expression is essential for maintaining cellular identity and function, and its disruption is a hallmark of cancer. While genetic and epigenetic changes are well-established drivers of transcriptomic variability in tumors, accumulating evidence indicates significant contributions from post-transcriptional mechanisms^1^. In breast cancer, where metastasis is the primary cause of mortality^2^, these changes are often driven by alterations in gene regulatory networks. Despite growing recognition of post-transcriptional regulation’s impact on cancer, the extent to which mechanisms such as mRNA stability contribute to gene expression variation across cancer transcriptomes remains unexplored^3,4^. Recent work by our group and others has underscored the importance of post-transcriptional mechanisms in cancer progression^5–7^, highlighting the need for a more systematic and quantitative assessment of these regulatory processes in breast cancer.

RNA stability is regulated through the combined action of cis-acting RNA sequence elements and trans-acting factors such as RNA-binding proteins (RBPs). On the cis side, specific sequence or structural motifs, often embedded within the transcript’s 3’ untranslated region (3’UTR), modulate transcript stability by providing direct binding sites^8–10^. On the trans side, RBPs recognize and bind these motifs, interacting with other RBPs and effector proteins to form regulatory complexes^11^. These regulatory modules then stabilize or destabilize targeted mRNA transcripts. Historically, cis and trans determinants have been studied separately, with interactions combined only post-hoc^8,12–14^. However, because multiple RBPs can recognize the same sequence element and vice versa, post-transcriptional regulation inherently involves many-to-many relationships. To better capture this complexity, we developed GreyHound, a computational framework that jointly integrates RNA sequence features and RBP expression data to predict transcript stability and identify candidate post-transcriptional regulatory networks. Unlike previous approaches, GreyHound enables simultaneous evaluation of both cis and trans determinants, providing a comprehensive view of RNA stability regulation. We applied GreyHound to RNA stability and expression profiling data from six breast cancer cell lines representing diverse breast cancer subtypes. Although transcript decay rates were broadly conserved across these cell lines, we observed notable variations contributing to cancer heterogeneity. Using GreyHound, we identified candidate cis–trans regulatory pairs that influenced transcript stability across these breast cancer models. The regulatory interaction between the RNA-binding protein RBMS3 (RNA Binding Motif Single Stranded Interacting Protein 3) and its associated sequence element emerged as the most significant regulatory axis.

Through loss- and gain-of-function studies, we confirmed RBMS3 as a key post-transcriptional regulator of mRNA stability. Consistently, reduced expression of RBMS3 was strongly associated with disease progression and aggressive clinical phenotypes in patient cohorts. Analysis of the RBMS3 target regulon revealed significant enrichment for metastasis-related biological pathways, prompting us to investigate metastatic potential downstream of this regulatory program. Overexpression of RBMS3 reduced metastatic potential and its knockdown enhanced metastatic lung colonization across independent xenograft models of breast cancer. Using a systematic in vivo CRISPR-interference screen, we identified TXNIP (Thioredoxin-interacting protein) as a critical downstream target of RBMS3; depletion of TXNIP resulted in similar phenotypes as RBMS3 loss-of-function phenotypes, and subsequent in vivo lung colonization experiments confirmed the epistatic relationship between RBMS3 and TXNIP. Collectively, our study uncovered a previously unknown metastasis-suppressive regulatory program mediated through post-transcriptional stabilization of a specific regulon, demonstrating the capability of our computational model, GreyHound, to reveal clinically relevant post-transcriptional regulatory networks in cancer.

## Results

### Heterogeneity in mRNA dynamics across cell line models of breast cancer

To systematically characterize RNA dynamics across different breast cancer models and subtypes, we performed metabolic RNA labeling using SLAM-seq (thiol(SH)-linked alkylation for the metabolic sequencing of RNA)^15^ on six breast cancer cell lines representing three major subtypes: luminal (ZR-75, MCF7), HER2-positive (MDA-MB-453), and triple-negative breast cancer (TNBC; HCC1806, MDA-MB-231, HCC38) (**Fig 1A**). As expected, transcription rates largely correlated with gene expression across these cell lines, explaining a substantial fraction of gene expression variation (**Fig 1B**). However, a significant portion of gene expression variability remained unexplained by transcriptional changes alone (**Fig 1C**), suggesting that additional regulatory mechanisms contribute to variations in mRNA abundance. We hypothesized that post-transcriptional control, particularly differential RNA decay rates, likely account for the remainder of this variation.

**Figure 1:**
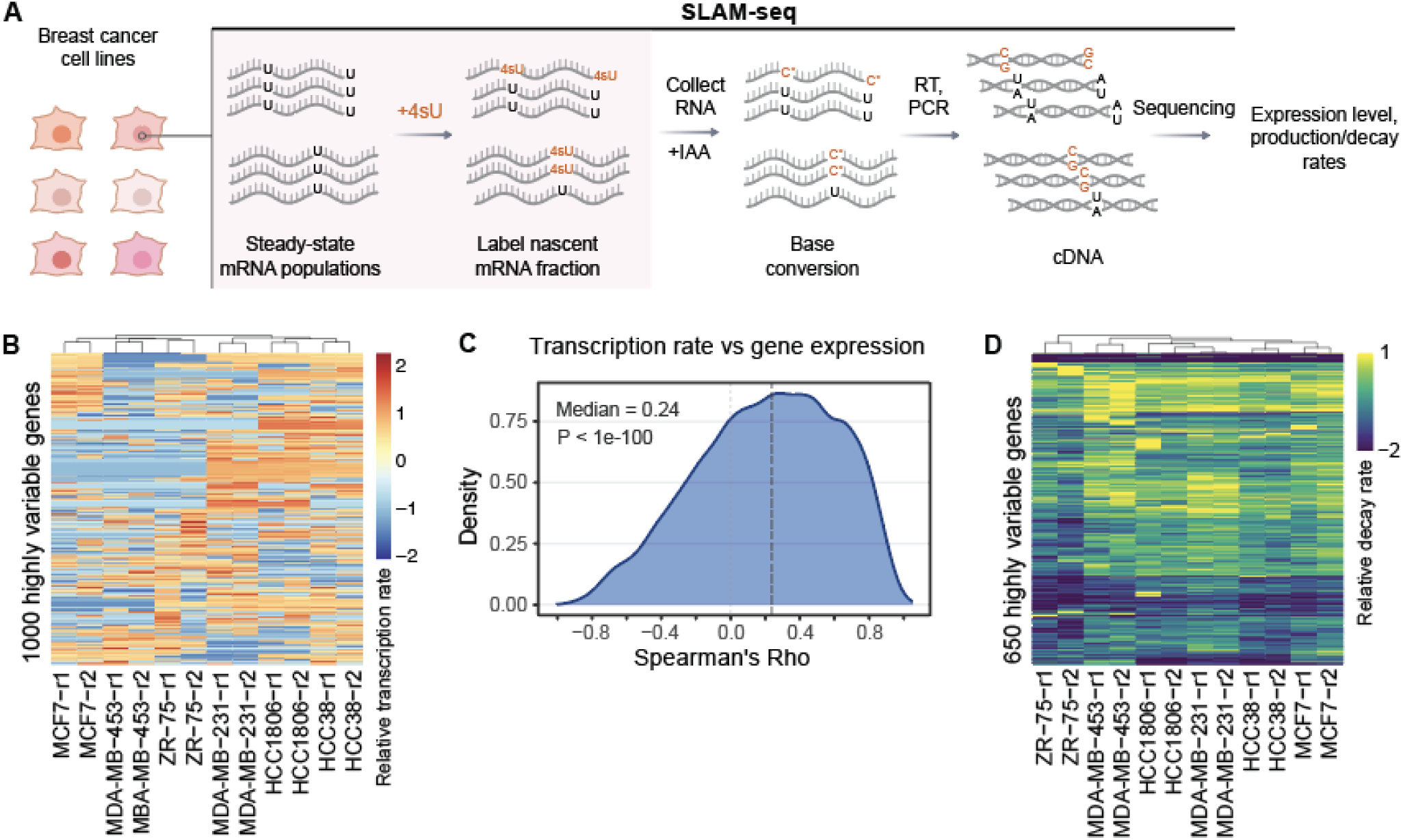
SLAM-seq reveals heterogeneity in mRNA transcription and stability in models of breast cancer. **(A)** Schematic of SLAM-seq showing how 4sU labeling causes T to C nucleotide conversions during library preparation and are detected during analysis of sequencing data to determine metabolic labeling rate. This in conjunction with expression data can be used to estimate mRNA decay in cell lines. **(B)** Normalized transcription rates based on observed C/T ratios from SLAM-seq; shown are the top thousand highly variable genes in this dataset. Note that the columns are clustered using hierarchical clustering and biological replicates group together. **(C)** A density plot showing the distribution of the Spearman correlations between SLAM-based transcription rates and expression for each gene across samples. **(D)** A heatmap of normalized mRNA decay rate estimates across cell lines and replicates for 650 highly variable genes.

We used gene expression values and metabolic labeling rates from SLAM-seq data to estimate mRNA decay rates in each sample. These decay rate measurements were highly reproducible across biological replicates (average Pearson R ≈ 0.9; **Fig. S1A**), validating the robustness of our profiling approach. While global mRNA decay patterns were broadly conserved across cell lines (average pairwise R = 0.59), we identified substantial heterogeneity in decay rates for hundreds of transcripts (**Fig. 1D**). Principal component analysis of these decay profiles revealed subtype-specific structures as well, with triple-negative breast cancer (TNBC) models clustering together (**Fig S1B**). Consistently, ANOVA revealed ∼100 genes with statistically significant cell line-specific decay patterns (P < 0.003, FDR < 0.25), reflecting meaningful post-transcriptional differences across breast cancer subtypes (**Fig S1C**).These findings motivated us to explore the underlying determinants of variability in RNA stability, which we describe in subsequent sections.

**Figure S1:**
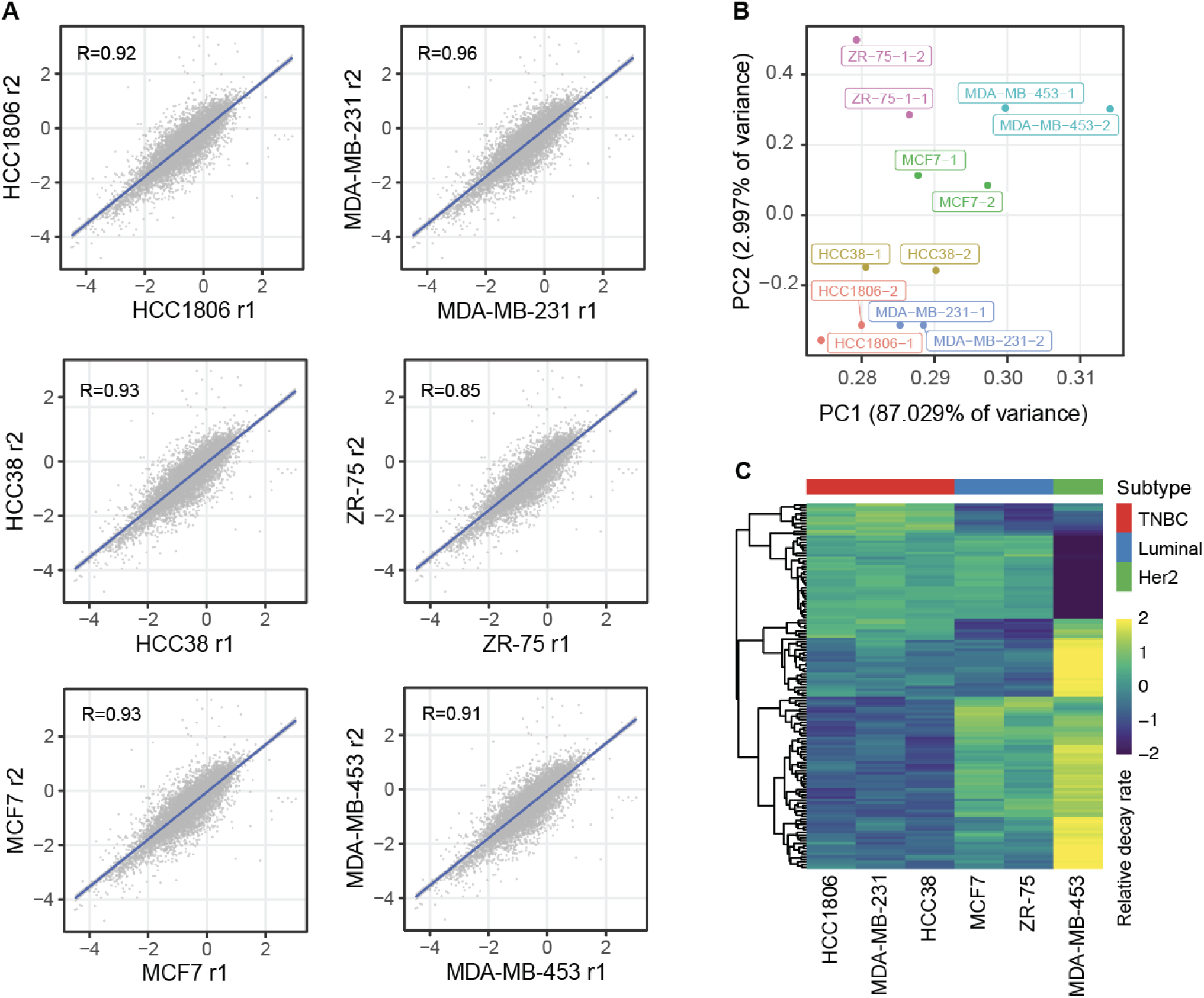
SLAM-based mRNA decay estimates across biological replicates, cell lines, and breast cancer subtypes. (**A**) Scatter plots demonstrating the correlation between mRNA decay rate estimates between biological replicates for each cell line. (**B**) PCA plots summarizing the mRNA decay rate measurements across samples. (**C**) A heatmap of normalized mRNA decay rate estimates across cell lines for 136 subtype-specific genes.

### Integrative modeling of mRNA stability reveals post-transcriptional heterogeneity in breast cancer

While mRNA decay rates are similar across breast cancer cell lines, there are also notable cell-state specific variations. In order to understand the role of post-transcriptional control that gives rise to (and may be influenced by) heterogeneity in breast cancer, we sought to reveal the *cis*-*trans* regulatory interactions that underlie the differential regulation of mRNA stability across breast cancer lines. To comprehensively capture the complexity of RNA stability regulation across diverse cellular contexts, we developed GreyHound, a deep learning framework that jointly integrates RNA sequence features and RNA-binding protein (RBP) expression data (**Fig 2A**). Unlike previous approaches, GreyHound simultaneously evaluates both *cis* and *trans* determinants of transcript stability, enabling the identification of context-dependent post-transcriptional regulatory interactions. The model architecture employs convolutional layers for sequence processing coupled with a pre-trained variational autoencoder for RBP expression encoding (**Fig S2A-B**). As shown in **Fig 2B**, given the expression profile of RBPs in a cell and the mRNA sequence of interest, GreyHound predicts the estimated decay rate for that mRNA within the cellular context defined by its RBP expression profile. Evaluating the performance of this model on a held-out test set showed a general agreement between the predicted and measured mRNA decay rates across pairs of genes and cell lines (R=0.62).

**Figure 2:**
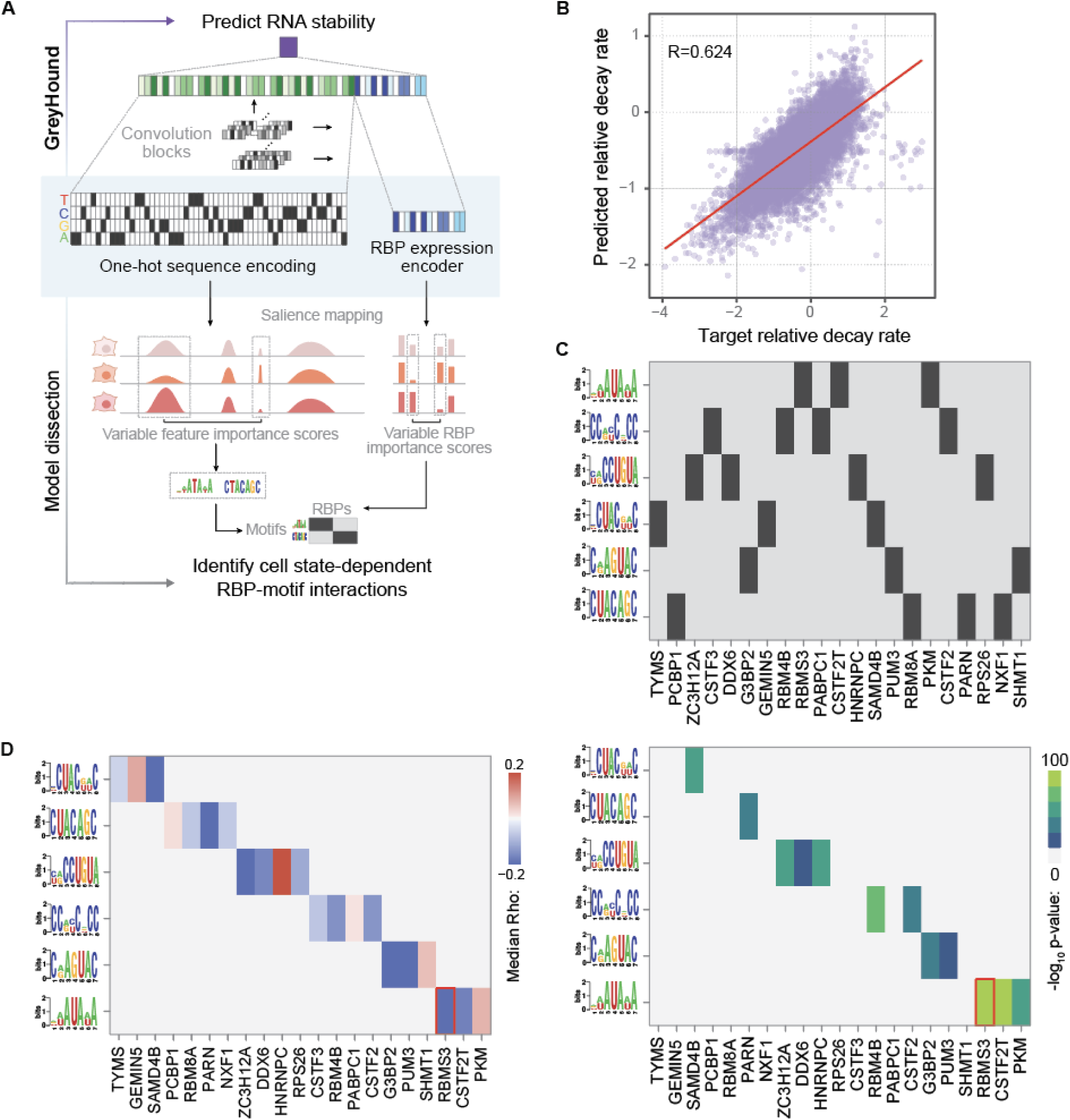
GreyHound reveals the *cis* and *trans* post-transcriptional regulatory programs in breast cancer cells. **(A)** Our strategy for measuring feature importance using model gradients at input. As denoted here, we specifically focused on features that show differential importance across cell lines. **(B)** Performance of our GreyHound model on a held-out test set. Each point represents the measurement and prediction for a given gene and a given cell line. **(C)** A map of putative RBP-motif interaction revealed by motif enrichment analysis among the sequences attributed to each RBP of interest. **(D)** Further evaluation of these interactions by analyzing the distribution of Spearman correlations between the expression of each RBP, and the decay rate measurements for its putative target transcripts based on motif occurrence. Shown are the median Spearman correlation for each nominated RBP-regulon pair (left) and its associated *p*-value calculated using Wilcoxon rank sum test (right). The interaction between RBMS3 and an AUA element is highlighted as the most significant interaction.

**Figure S2:**
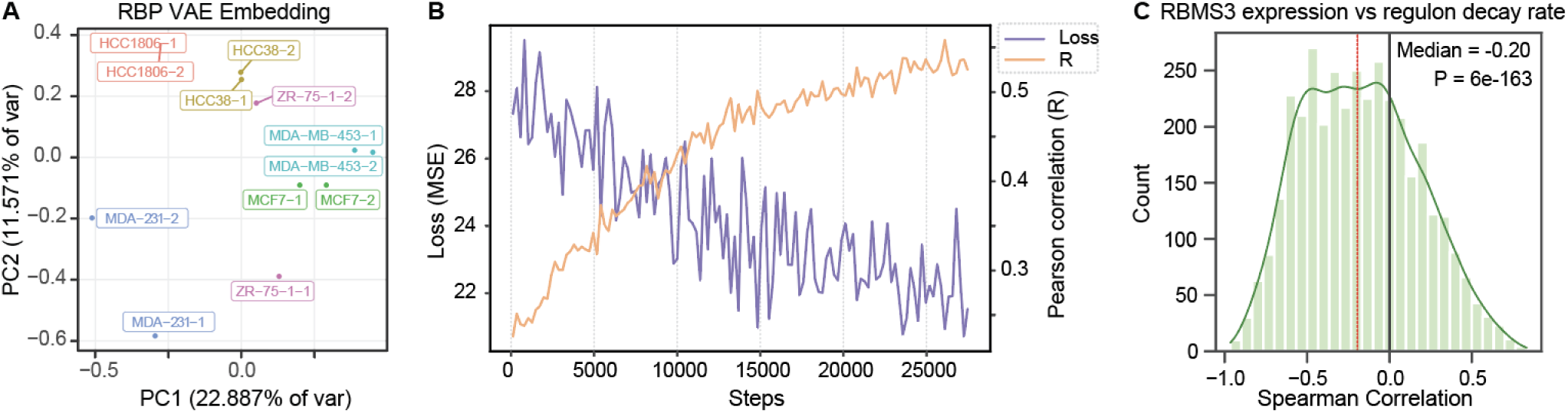
Nomination of RBMS3 as a regulator of differential mRNA stability across breast cancer cell lines. (**A**) PCA plot of the embedding learned for each sample using the expression of RBPs. This is to highlight that RBP expression can largely capture cell states. (**B**) The training loss and Pearson correlation curves for the model. The model was stopped when the validation R had largely plateaued. (**C**) The distribution of Spearman correlation coefficient between RBMS3 expression and decay rates for its putative target transcripts. The median and associated *p*-value (Wilcoxon test) are also shown.

To decipher the cell-state dependent regulatory logic learned by the model, we implemented a comprehensive model dissection approach that combines gradient-based saliency analysis with rigorous statistical validation (**Fig 2A**). For each gene-cell line pair, we estimated feature importance across the sequence as well as RBPs by calculating gradient at input (i.e. akin to saliency). For both heads of the model, we specifically focused on features whose importance shows variation across cell states. For sequence analysis, we identified regions of variable importance across cell lines by modeling the mean-variance relationship and identifying deviations from the expected variance. We used a smoothened coefficient of variation (CV) across each sequence to mark highly variable regions (top 10% of CV). These regions were merged if they occurred within 20 nucleotides of each other, defining key regulatory regions relevant to cell-state dependent predictions of the model. In parallel, for RBP inputs, we similarly identified proteins with consistently highly variable importance scores across conditions. To pinpoint the underlying *cis*-regulatory elements over-represented across the discovered highly variable regions, we used FIRE (Finding Informative Regulatory Elements) to perform de novo motif discovery^13^. To link these sequence motifs to the selected RBPs, we performed a pair-wise enrichment analysis for each motif among the sequences for which the RBP was identified as being differentially salient (**Fig 2C**). The statistical significance of these RBP-motif interactions was further validated through Spearman correlation analysis between RBP expression and target transcript stability (**Fig 2D**). The most significant interaction within this predicted network of post-transcriptional regulatory interactions was between RBMS3 and an AUA-rich sequence element, highlighted by red boxes in **Figure 2D**. As shown in **Fig S2C**, the decay rates measured across cell lines for the RBMS3 regulon, defined as the set of transcripts that carry the identified AUA elements, were on average anti-correlated with the expression of this RBP in these lines. Our RBP-motif interaction map described here provides a quantitative framework for understanding post-transcriptional regulation of RNA stability across cell states, and nominates RBMS3 as a potential regulator of mRNA stability in the context of breast cancer.

### RBMS3 acts as a post-transcriptional regulator of mRNA stability in breast cancer cells

Following the identification of RBMS3 as a potential regulator of mRNA stability in breast cancer by our GreyHound model, we next sought to investigate the functional impact of RBMS3 expression on its target regulon. To this end, RBMS3 was silenced in the MDA-MB-231 breast cancer cell line, which exhibits relatively high endogenous RBMS3 expression compared to other breast cancer cell lines, using two independent short hairpin RNAs (shRNAs). Successful knockdown of RBMS3 was confirmed prior to an RNA-seq analysis to assess gene expression changes resulting from RBMS3 depletion (**Fig S3A**). Motif enrichment analysis using FIRE^13^ revealed a strong and highly significant over-representation of the AUA motif within the 3’UTR of transcripts that were downregulated in RBMS3 knockdown cells (**Fig 3A**). To ensure that this observation was not an artifact of motif selection, we utilized RBMS3 binding scores from DeepBind (D00138.001) to identify transcripts that are likely targets of RBMS3 based on its binding preferences learned from RNA compete assays^16^. As shown in **Fig S3B**, the DeepBind-predicted RBMS3 regulon were also significantly downregulated upon RBMS3 knockdown, supporting the hypothesis that RBMS3 affects the stability of its target transcripts.

**Figure 3:**
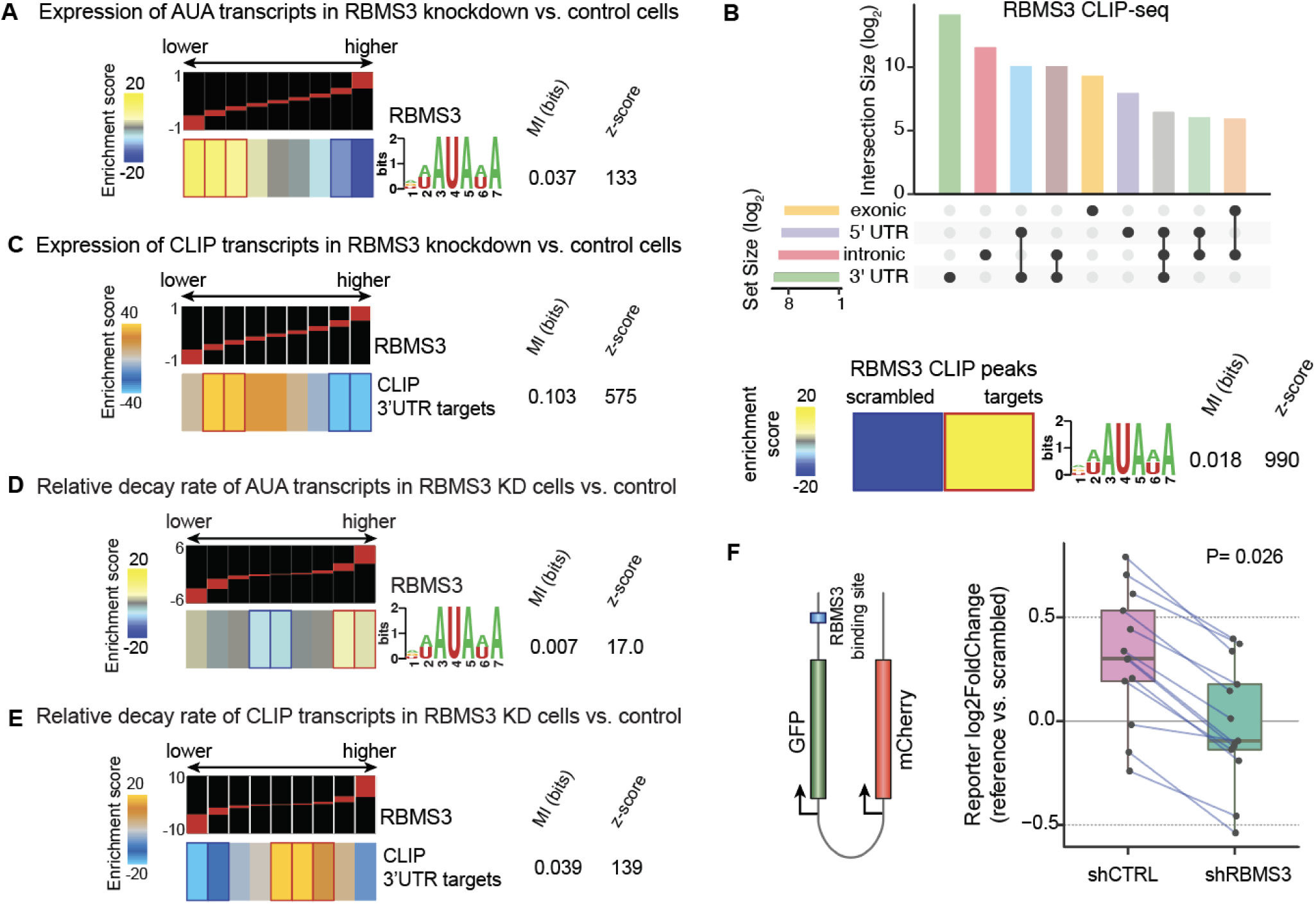
RBMS3 binding impacts target regulon expression and stability in breast cancer cells. (**A**) Enrichment and depletion patterns of AUA element containing transcript’s expression level in breast cancer cell line MDA-MB-231 with RBMS3 knocked down versus control cells. (**B**) RBMS3 binding site locations identified by CLIP-seq (top). Enrichment of the AUA motif among RBMS3 CLIP-seq binding sites (bottom). (**C**) Enrichment and depletion patterns of RBMS3 target regulon expression in breast cancer cell line MDA-MB-231 with RBMS3 knocked down versus control cells. (**D**) Enrichment and depletion patterns of RBMS3 target AUA motif-containing transcripts relative RNA decay rate in breast cancer cell line MDA-MB-231 with RBMS3 knocked down versus control cells. (**E**) Enrichment and depletion patterns of RBMS3 target regulon relative RNA decay rate in breast cancer cell line MDA-MB-231 with RBMS3 knocked down versus control cells. (A, C-E) Shown are enrichment patterns along with calculated MI and associated Z-score. See ref ^13^ for details. (**F**) Dual reporter system that expresses mCherry and GFP driven by a bi-directional CMV lentiviral promoter. RBMS3 binding recognition sites are inserted into the 3’UTR of GFP (Left). Expression analysis of paired target sequence to scrambled control sequence in MDA-MB-231 RBMS3 knockdown versus control cells (Right).

**Figure S3:**
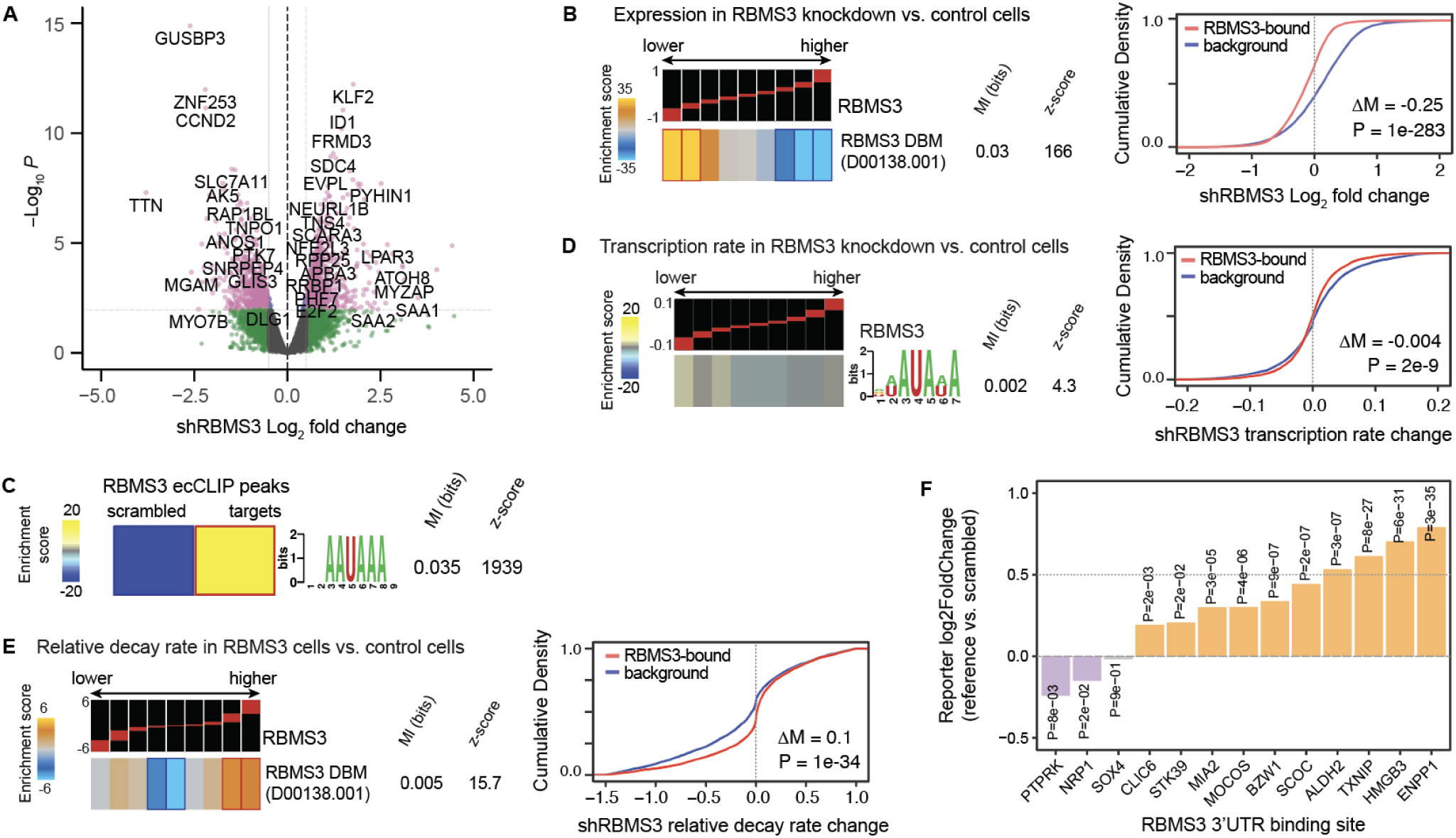
RBMS3 expression impacts putative and experimental target regulon expression and stability. (**A**) Volcano plot of differential gene expression in MDA-MB-231 RBMS3 knockdown cells versus control cells(n=4). (**B**) Enrichment and depletion patterns of the DeepBind RBMS3 regulon shows RBMS3 target gene enrichment among the genes that are downregulated in RBMS3 knockdown cells. (**C**) De novo motif discovery in RBMS3 binding sites, with dinucleotide-invariant scrambled sequences as background, identified an AUA motif, similar to the one we identified based on our analysis of GreyHound’s feature importance scores. (**D**) Enrichment and depletion patterns of the transcription rates for transcripts containing the AUA sequence element in MDA-MB-231 RBMS3 knockdown versus control cells. (**E**) Enrichment and depletion patterns of the relative decay rates of DeepBind RBMS3 regulon in MDA-MB-231 RBMS3 knockdown versus control cells. (B, D-E) Shown are enrichment patterns along with calculated mutual information (MI) and associated Z-score (see ref ^13^ for details), in addition to the cumulative density. (**F**) The contribution of each cloned target binding sequence on GFP expression by comparing their relative abundances in GFP mRNA to that of their scrambled controls. We performed this analysis in both control and RBMS3 knockdown cells, and used a paired Wilcoxon signed rank test to report a *p*-value.

To further validate our findings regarding RBMS3 and its putative regulon, we experimentally identified the RBMS3-bound target transcripts using a modified version of CLIP-seq (cross-linking and immunoprecipitation sequencing)^17,18^. Since reliable antibodies for immunoprecipitation of endogenous RBMS3 under CLIP conditions were not available, we expressed FLAG-tagged RBMS3 in MDA-MB-231 cells to perform transcriptome-wide mapping of its binding sites. Consistent with our computational predictions, we observed (i) extensive RBMS3 binding within 3’UTRs across thousands of sites, and (ii) a significant enrichment of the AUA motif previously identified through GreyHound analysis (**Fig 3B**). De novo motif discovery among the RBMS3 CLIP binding sites also identified an AUA element, providing evidence for *in vivo* binding of RBMS3 to this motif (**Fig S3C**). Furthermore, when we repeated our analysis from the previous section with this experimentally defined set of RBMS3-bound targets instead of the motif-based putative regulon, we found that these transcripts were also significantly downregulated in RBMS3 knockdown cells (**Fig 3C**). These findings provide strong evidence that RBMS3 binds and stabilizes its target transcripts through interactions with AUA-containing elements in their 3’UTRs.

In order to ensure that the observed reduction in transcript abundance in response to RBMS3 silencing is due to post-transcriptional changes in mRNA stability and not transcriptional modulations, we performed SLAM-seq^15^ in MDA-MB-231 cells following RBMS3 knockdown. Consistent with our previous observations, transcripts exhibiting higher decay rates upon RBMS3 depletion were significantly enriched for the 3’UTR-AUA motif (**Fig 3D**). However, the AUA motif did not significantly affect differential transcriptional measurements: transcripts containing this element were neither enriched nor depleted in transcriptional output (**Fig S3D**), indicating that RBMS3’s primary mode of action is post-transcriptional rather than transcriptional. To confirm the robustness of these findings across different target definitions, we replicated the analysis using both the DeepBind predicted regulon (**Fig S3E**) and the RBMS3 CLIP-derived target set (**Fig 3E**). In each case, transcripts bearing the AUA motif exhibited significantly increased decay rates upon RBMS3 depletion.

To directly test whether RBMS3 binding sites are sufficient and required for RBMS3-mediated regulation of mRNA stability, we employed a dual-reporter system previously established for investigating post-transcriptional regulatory elements in living cells^9,19^. From a set of high-confidence RBMS3-bound 3′UTRs (described in later sections), we selected representative binding sites for functional validation. These sequences were cloned downstream of a GFP coding region in our dual-reporter vector, along with matched scrambled control sequences that preserved di-nucleotide composition but disrupted motif content.

This reporter library was transduced into MDA-MB-231 control and RBMS3 knockdown cells, and targeted RNA-seq was used to quantify reporter expression as a readout of *cis*-element activity and its dependence on RBMS3. We made two key observations consistent with our molecular model of RBMS3 action: (1) in most cases, the presence of the RBMS3 binding site increased reporter mRNA levels relative to scrambled controls (**Fig S3F**), and (2) this effect was absent in RBMS3-depleted cells, indicating that the observed stabilization was dependent on RBMS3 expression (**Fig 3F**). Together, these findings demonstrate that direct RBMS3-RNA interactions are both necessary and sufficient to confer increased transcript stability.

### RBMS3 and its downstream target regulon are associated with breast cancer progression

As we have demonstrated in the preceding sections, RBMS3 modulates the stability of its target mRNA transcripts. Next, we wanted to delineate which downstream molecular and cellular pathways are impacted by RBMS3. Therefore, we conducted a series of gene-set enrichment analyses. First, we employed Appyters^20^ to identify pathways that are enriched among the transcripts bound by RBMS3 based on CLIP-seq annotations. As shown in **Fig 4A**, we identified VEGF and TGF-β pathways as the top associated gene-sets. Parallel analyses of gene and protein expression changes in response to RBMS3 knockdown similarly identified gene-sets associated with breast cancer progression and metastasis (**Fig S4A-B**). These results raised the possibility that RBMS3 plays a role in breast cancer progression. Specifically, our findings indicate that RBMS3 silencing is associated with increased expression of oncogenic and pro-metastatic pathways.

**Figure 4:**
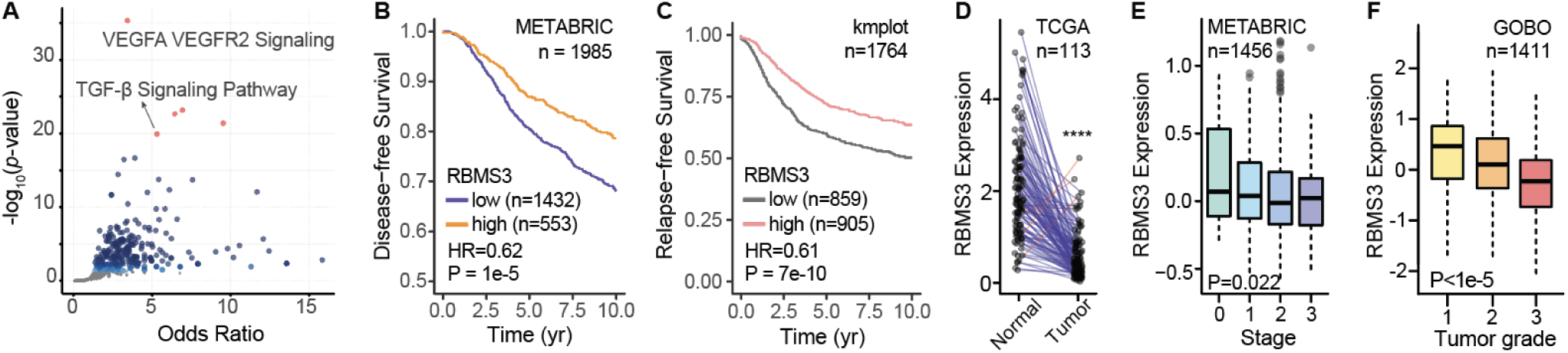
RBMS3 is a predictor of breast cancer progression. (**A**) Gene-set enrichment analysis using Appyters^20^ to identify pathways that are enriched among the transcripts bound by RBMS3 based on CLIP-seq annotations. VEGF and TGF-β pathways were top associated gene-sets. (**B**) Analysis of disease-free survival in the METABRIC cohort^21^ relative to RBMS3 expression. (**C**) Meta-analysis of relapse-free survival in smaller published cohorts^22^ relative to RBMS3 expression. (B-C) The Mantel-cox test was used to measure significance. Low RBMS3 expression is indicative of poor prognosis for disease-free and relapse-free survival in breast cancer patients. (**D**) Analysis of paired normal and breast cancer samples in the TCGA dataset revealed an almost universal reduction in RBMS3 expression in tumor tissue. Paired Mann-Whitney test was used. (**E**) RBMS3 expression in METABRIC cohort^21^ by tumor stage (0-1: early, 2-3: late). Mann-Whitney U test was used for statistical comparison. (**F**) RBMS3 expression across 1411 tumor samples stratified based on their annotated grades (GOBO meta-analysis)^24^. ANOVA was used to calculate the associated p-value.

**Figure S4:**
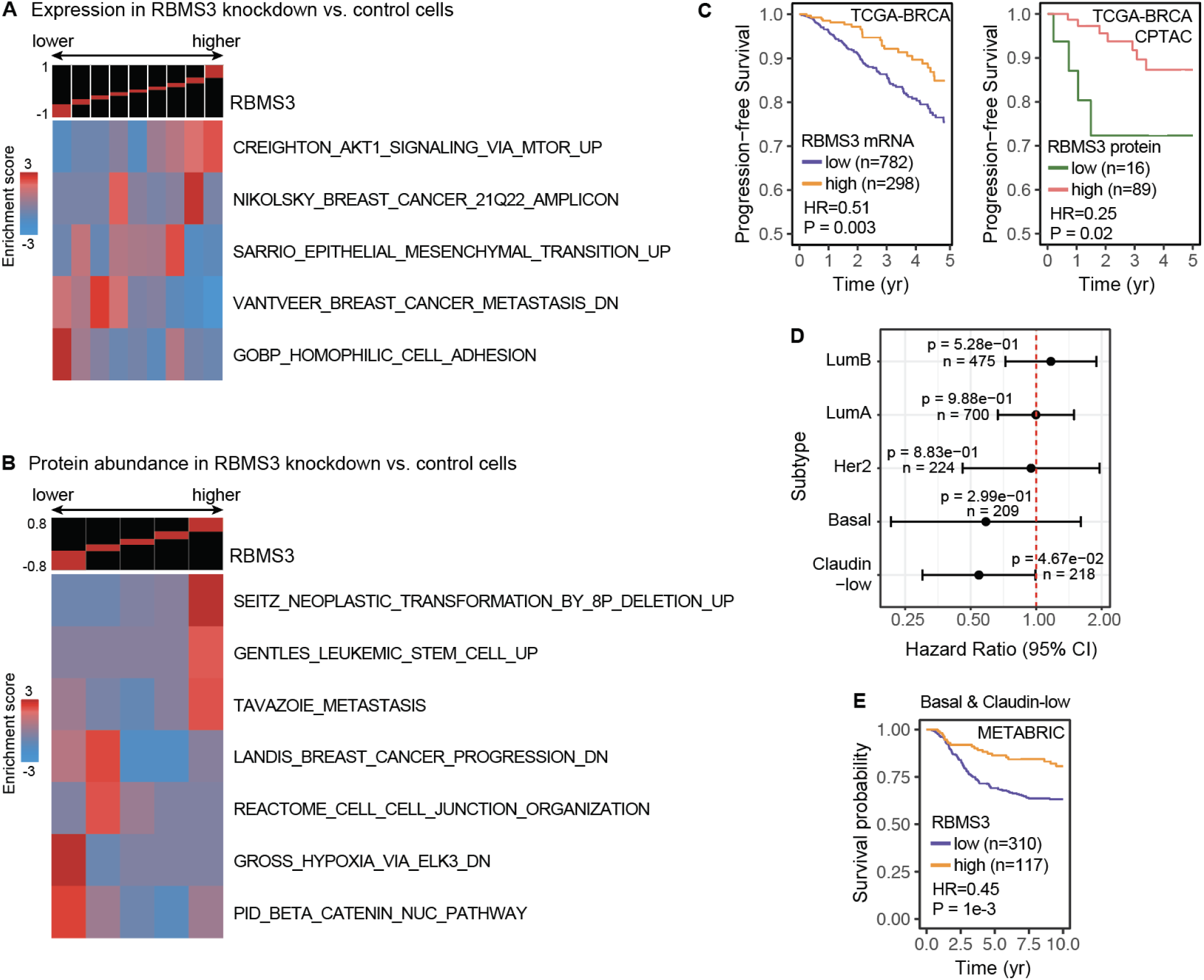
RBMS3 is a predictor of breast cancer progression. (**A**) Cancer pathway associated gene expression enrichment analysis in RBMS3 knockdown versus control cells. (**B**) Cancer pathway associated protein abundance enrichment analysis in RBMS3 knockdown versus control cells. (**C**) Analysis of progression-free survival in the TCGA-BRCA and TCGA-BRCA CPTAC cohorts relative to RBMS3 expression. TCGA-BRCA (The Cancer Genome Atlas – Breast Invasive Carcinoma Cohort) and TCGA-BRCA-CPTAC (Clinical Proteomic Tumor Analysis Consortium) data sets were downloaded from the genome data commons. (**D**) Hazards ratio 95% confidence interval of subtypes in the METABRIC cohort^21^. (**E**) Analysis of survival probability in the Basal and Claudin-low subtypes of the METABRIC cohort^21^.

We next sought to evaluate the association between RBMS3 expression and clinical outcomes in breast cancer using patient data from several independent breast cancer datasets. Analysis of the METABRIC dataset^21^, containing gene expression profiles for ∼2000 patients, revealed that diminished RBMS3 levels were indicative of poor clinical outcomes (**Fig 4B**). Data from a meta-analysis of existing breast cancer datasets^22^ likewise revealed a negative association between RBMS3 expression and relapse-free survival (**Fig 4C**). A negative association of RBMS3 expression and progression-free survival was also observed in the analysis of both RBMS3 mRNA and protein levels in the TCGA-BRCA and TCGA-BRCA-CPTAC breast cancer cohorts (**Fig S4C**). Additionally, RBMS3 expression was consistently reduced in higher grade and later stage tumors across several independent breast cancer cohorts (**Fig 4D-F**). These findings are consistent with prior research that analyzed RBMS3 expression in patient data from the TCGA dataset and similarly reported that reduced RBMS3 expression is associated with poor clinical outcomes^23^. Taken together, these results demonstrate a significant association between RBMS3 loss and adverse clinical outcomes, highlighting its potential utility as a prognostic biomarker in breast cancer.

Finally, we observed that the negative association observed between RBMS3 expression and survival was largely attributed to the Basal and Claudin-low subtypes in the METABRIC cohort^21^ (**Fig S4D-E**). Based on this observation—and given the known increased metastatic potential of these subtypes—we focused subsequent functional and phenotypic analyses specifically on the triple-negative models of breast cancer.

### RBMS3 modulation impacts breast cancer metastasis and invasion

To establish a causal link between RBMS3 silencing and higher metastatic potential, we performed lung colonization assays in immunodeficient NSG (NOD scid gamma) mice. We injected MDA-MB-231 cells with RBMS3 knocked down (two independent shRNAs) or control cells via tail-vein and measured metastatic burden in the lungs over time using *in vivo* imaging (**Fig 5A**). Consistent with our earlier findings, mice injected with RBMS3 knockdown cells exhibited significantly greater metastatic lung colonization compared to those injected with control cells. We also established that this increase is not attributable to higher proliferation rate in these cells (**Fig S5A**).

**Figure 5:**
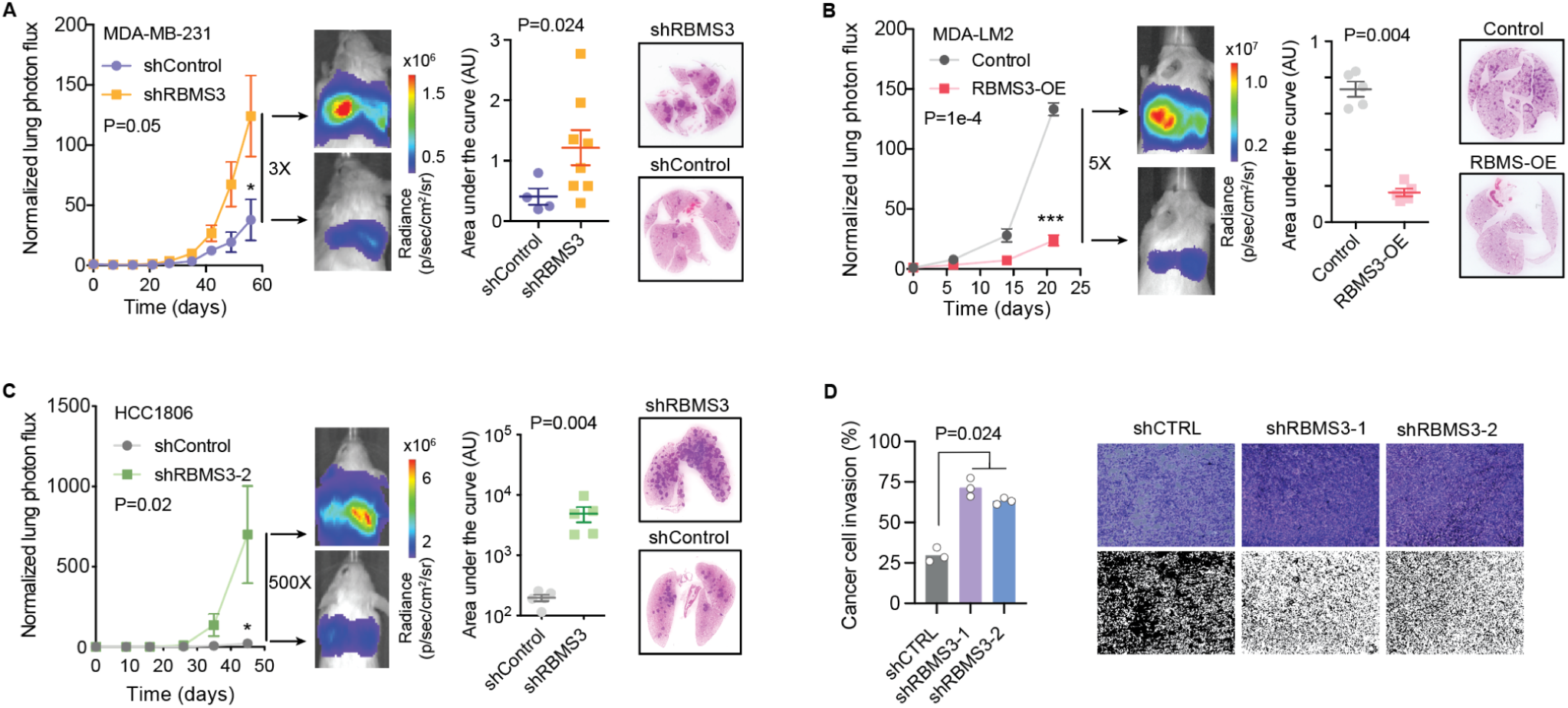
Loss of RBMS3 increases metastatic lung colonization and invasion phenotypes. (**A-C**) Lung colonization assay for control and either RBMS3 knockdown or RBMS3 overexpression. Luciferase-labeled cells were injected via tail-vein and their metastatic growth in the lungs was measured over time. The magnitude of the signal, shown here as a heat map for a representative mouse from each cohort, reflects the metastatic burden. Also included are area under the curve (AUC) of log-normalized signal in the lungs of mice over 25 days and H&E stained lung sections for exemplary mice. (**A**) Lung colonization assays of control (N=4) and RBMS3 knockdown MDA-MB-231 cells (Total N=8). (**B**) Lung colonization assays for control (N=5) and RBMS3 over expression MDA-LM2 cells (N=5). (**C**) Lung colonization assays for control (N=5) and RBMS3 knockdown HCC1806 cells (N=5). (**D**) Invasion assays for control and RBMS3 knockdown MDA-MB-231 cells. Images are representative (i.e. median) in each group. In processed images, black is empty space and white is cells. The graph shows fraction that is white (i.e. % area covered) and the Mann-Whitney U test was used for statistical comparison.

**Figure S5:**
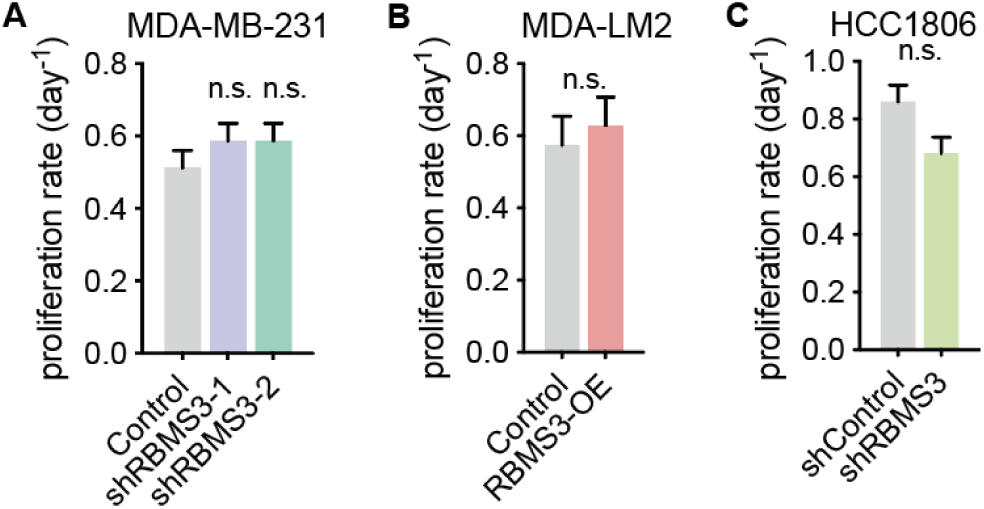
RBMS3 expression does not affect proliferation. (**A**) *In vitro* proliferation rate in RBMS3 knockdown and control cells (two independent hairpins in MDA-MB-231 cells; N = 3). (**B**) *In vitro* proliferation rate in RBMS3 over expression and control cells (in MDA-MB-231 cells; N = 3). (**C**) *In vitro* proliferation rate in RBMS3 knockdown and control cells (best performing hairpin was used in HCC1806 cells; N = 3).

To complement this loss-of-function approach, we performed a gain-of-function experiment, by over-expressing RBMS3 in the MDA-LM2 cell line, a highly metastatic derivative of MDA-MB-231^25^. Elevated RBMS3 expression resulted in a marked reduction in lung colonization (**Fig 5B**) again independent of the proliferation rates of these cells (**Fig S5B**). To ensure that our findings are not limited to the MDA-MB-231 background, we also included the HCC1806 cell line as an independent model of claudin-low breast cancer^26^. Consistent with our prior finding, shRNA-mediated silencing of RBMS3 led to a substantial and significant increase in metastatic lung colonization in xenografted NSG mice with no changes in proliferation rates (**Fig 5C & S5C**).

Given the absence of proliferation differences, and considering that metastasis is also influenced by cancer cell invasiveness, we next assessed whether RBMS3 affects invasive behavior using transwell invasion assays. RBMS3 knockdown in MDA-MB-231 cells led to a significant increase in invasive capacity (**Fig 5D**) indicating that RBMS3 plays an important role in regulating this key hallmark of breast cancer metastasis^2^. Collectively, these results establish RBMS3 as a novel suppressor of metastasis and invasion in claudin-low/basal breast cancers.

### The RBMS3-TXNIP regulatory axis suppresses breast cancer metastasis

Our results so far indicate that (1) RBMS3 functions as a regulator of mRNA stability, (2) it is associated with breast cancer progression in basal-like/claudin-low subtypes, (3) RBMS3 expression suppresses metastatic lung colonization in cell line derived xenograft models. However, to confirm that RBMS3’s metastasis-suppressive role is connected to its function as a post-transcriptional regulator and to better understand the molecular mechanisms downstream of RBMS3, we sought to identify its target transcripts that contribute to metastasis. To select these candidate genes, we focused on genes that show direct RBMS3 binding in their 3’UTRs. We observed that for these genes, the changes in gene expression in response to RBMS3 silencing is overall correlated at the mRNA and protein levels (Spearman correlation of 0.2, *P*=2e-19). Among these genes, we selected those that were significantly downregulated in both datasets (logFC<-0.25 and *p*-value<0.05). We also selected transcripts with no reduction in transcription rate based on our SLAM-seq data. Finally, we also performed survival analyses in the METABRIC dataset^21^ and selected those genes that are associated with better survival outcomes. These criteria resulted in the selection of 13 genes.

Due to this set being too large for individual functional validation in xenograft models, we instead opted to conduct an *in vivo* pooled CRISPR-interference screen in NSG mice. We used the dual-guide system, where two guides are simultaneously cloned for each target gene to ensure effective knockdown^27^. MDA-MB-231 cells were transduced with the guide library that included 10 non-targeting controls and these cells were then injected into mice tail vein (N=3). In parallel, we also collected *in vitro* proliferated samples to account for potential changes in proliferation rates and to focus on the *in vivo* metastasis phenotype observed for RBMS3. Once lungs were colonized and metastatic foci were formed, we extracted the lungs of the mice, and isolated genomic DNA (**Fig 6A**). We then used targeted PCR for dual guide libraries and high-throughput sequencing to quantify the abundance of cells expressing each guide within the *in vivo* and *in vitro* grown cells. Among the 13 candidates, TXNIP emerged as showing a significant lung colonization phenotype, without impact on proliferation (**Fig 6B**). This phenotype mirrors that of RBMS3 knockdown, suggesting that TXNIP may in fact be a downstream effector of RBMS3 in suppressing metastasis.

**Figure 6:**
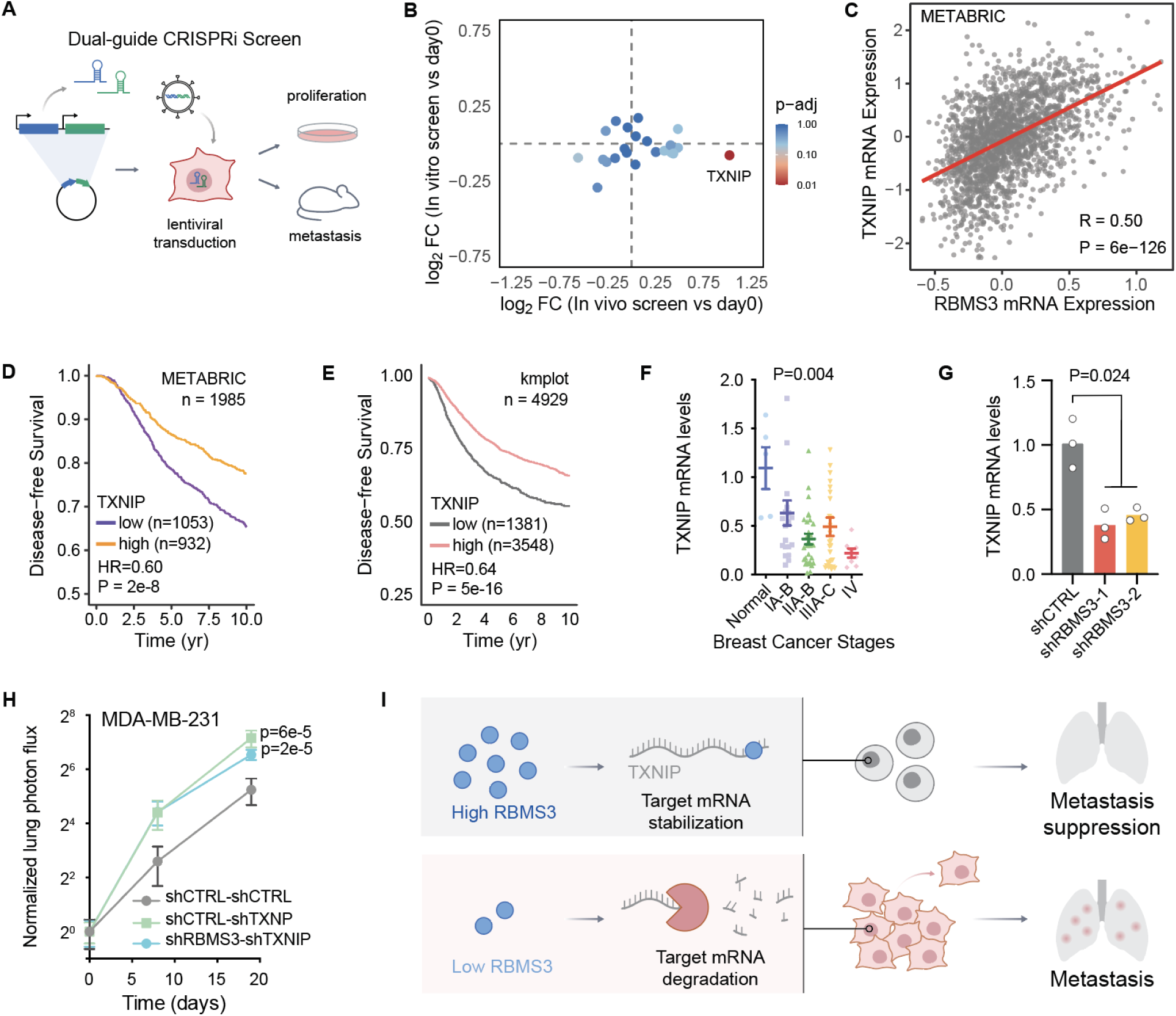
TXNIP knockdown shares phenotype with RBMS3 and acts as a tumor suppressor. **(A)** Schematic of the dual-guide CRISPR-interference screen. (**B**) Analysis of CRISPRi screen comparing the abundance of cells expressing each guide between *in vivo* and *in vitro* grown cells. TXNIP was observed to be present at high levels *in vivo* and low levels *in vitro* indicating high metastasis but low proliferation. (**C**) Comparative analysis of TXNIP to RBMS3 expression in the METABRIC cohort ^21^. (**D**) Analysis of disease-free survival in the METABRIC cohort^21^ relative to TXNIP expression. (**E**) Meta-analysis of relapse-free survival in smaller published cohorts^22^ relative to TXNIP expression. (D-E) The Mantel-cox test was used to measure significance. Low TXNIP expression is indicative of poor prognosis for disease-free survival in breast cancer patients. (**F**) The expression levels of TXNIP in 90 tumor samples at different stages of breast cancer was measured by qPCR. (**G**) The expression levels of TXNIP in RBMS3 knockdown MDA-MB-231 cells and control cells was measured by qPCR. (**H**) Lung colonization assays of control (N=5), TXNIP knockdown (N=5), and RBMS3-TXNIP double knockdown (N=5) in MDA-MB-231 cells. Luciferase-labeled cells were injected via tail-vein and their metastatic growth in the lungs was measured over time. (**I**) Molecular mechanism of RBMS3 metastasis suppression through post-transcriptional regulatory action.

Consistent with this model, we observed a significant and positive correlation between RBMS3 and TXNIP expression across multiple datasets, namely METABRIC^21^, TCGA, and MBCProject (**Fig 6C** and **Fig S6A-B**). Similarly, we also observed that lower expression of TXNIP is also strongly associated with poor outcomes (**Fig 6D-E**). To independently validate this, we measured TXNIP expression in a cohort of 90 breast tumor samples across different disease stages and found a significant association between TXNIP silencing and breast cancer progression (**Fig 6F**).

**Figure S6:**
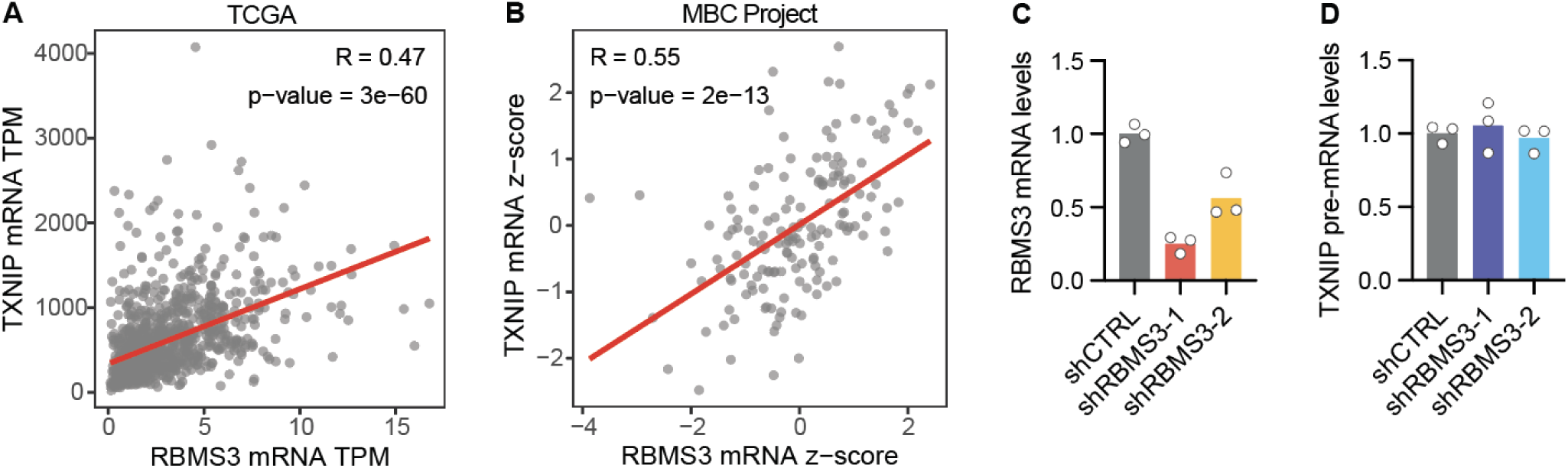
RBMS3 modulates TXNIP expression in a post-transcriptional manner. **(A-B)** Comparative analysis of TXNIP to RBMS3 expression in (A) TCGA and (B) MBCProject datasets (**C**) Expression levels of RBMS3 as measured by qPCR in RBMS3 knockdown and control cells. Sufficient knockdown of RBMS3 is seen. (**D**) Expression levels of TXNIP pre-mRNA in RBMS3 knockdown and control cells as measured by qPCR.

To confirm that TXNIP acts as a downstream target of RBMS3, we measured its pre-mRNA levels in RBMS3 knockdown cells using qPCR. Consistent with our previous observation of a ∼2-fold reduction in mRNA (P=1e-3) and protein levels (P=0.047) of TXNIP in RBMS3 knockdown cells, silencing RBMS3 resulted in a significant reduction in TXNIP mRNA levels (**Fig S6C** and **6G**). However, we observed no change in TXNIP pre-mRNA levels upon RBMS3 knockdown (**Fig S6D**), indicating that RBMS3 does not influence TXNIP transcription rates.

To further investigate the functional relationship between RBMS3 and TXNIP *in vivo*, we performed an epistasis experiment using MDA-MB-231 cells engineered to express a dual shRNA cassette targeting both RBMS3 and TXNIP. As shown in **Fig 6H**, knockdown of TXNIP alone significantly increased metastatic lung colonization relative to control cells, consistent with its role as a metastasis suppressor. Co-silencing of RBMS3 and TXNIP also enhanced metastatic colonization relative to controls. However, the extent of colonization in the double knockdown condition was not greater than that observed with TXNIP knockdown alone (**Fig 6H**). If RBMS3 and TXNIP acted through independent pathways, we would expect the double knockdown to yield an additive or synergistic increase in metastasis. Instead, the lack of an additive effect supports a model in which RBMS3 acts upstream to promote TXNIP expression.

## Discussion

Our study identifies the RBMS3-TXNIP post-transcriptional regulatory axis as a key suppressor of breast cancer metastasis. We demonstrate that RBMS3 binds and stabilizes the TXNIP transcript, and that loss of RBMS3 reduces TXNIP levels, leading to enhanced metastatic colonization without affecting proliferation (**Fig 6I**). This highlights a molecular mechanism in which RNA-binding proteins can exert metastasis-suppressive effects through regulation of transcript stability.

TXNIP has been previously implicated as a tumor suppressor in several cancer types^28^, including breast cancer^29,30^, though its regulation at the post-transcriptional level remains poorly understood. We show that RBMS3 is a direct, positive regulator of TXNIP mRNA stability, providing new mechanistic insight into how TXNIP is controlled. Notably, epistasis experiments confirmed that RBMS3 requires TXNIP to mediate its anti-metastatic effects, reinforcing the functional relevance of this regulatory axis.

The discovery of this pathway was enabled by our development of GreyHound, a computational framework that integrates RNA sequence and RBP expression data to infer regulators of transcript stability. GreyHound analysis pinpointed RBMS3 as a candidate stabilizer of AUA-containing transcripts. Experimental validation using CLIP-seq, dual reporter assays, and in vivo metastasis models confirmed that the RBMS3-target mRNA interaction, including with TXNIP, is required and sufficient for the observed regulatory effects.

This work expands the biological roles of RBMS3 beyond its prior characterization as a c-MYC–related RBP^31^. Our data indicate it acts as a broad regulator of mRNA stability with a particular role in constraining metastatic progression. These findings align with recent studies proposing tumor-suppressive roles for RBMS3^23^ and position it within the growing landscape of post-transcriptional regulators that influence metastasis.

More broadly, our approach provides a generalizable framework for uncovering regulatory pathways driven by mRNA stability. As transcriptome-wide mRNA stability measurements become more accessible, computational frameworks like GreyHound will enable systematic discovery of functional RNA-protein interactions across diverse disease contexts.

## Acknowledgements

We thank Brian Plosky and Chiara Ricci-Tam for reading earlier versions of this manuscript and helping us with the content. H.K. was a Zena Werb Scholar and was supported by T32CA108462. T.M. was supported by an MD fellowship from the Boehringer Ingelheim Fonds. Sequencing was performed at the UCSF Center for Advanced Technology (CAT), supported by UCSF PBBR, RRP IMIA, and NIH 1S10OD028511-01 grants. H.G. is a Core Investigator at Arc Institute, and his research is supported by Arc. H.G. is also an Era of Hope Scholar (W81XWH-2210121) and supported by grants from NCI (R01CA240984 and R01CA244634).

## Data and Code Availability

## Methods

### Genome-wide mRNA decay rate measurements across breast cancer cell lines

#### 4sU#metabolic labeling of mRNA

One day before the SLAM-seq procedure^15^, all cell lines were plated in biological replicates at 1×10^5^ cells/well on a 24-well-plate. For 4sU labeling, we incubated ZR-75-1, MDA-MB-453, HCC1806, MDA-MB-231, HCC38, and MCF7, as well as RBMS3 knockdown and control MDA-MB-231 cells in their respective culture media supplemented with 4-thiouridine (4sU, 100 µM) for 4 hours. After pulse labeling, the cells were washed twice with PBS and harvested directly in TRIzol Reagent (Invitrogen, 15596018). Total RNA was isolated and Iodoacetamide labeling was performed on all isolated RNA (input range 500 ng to 3 μg) using the SLAMseq Explorer and Kinetics Kits (Lexogen, 061), following the manufacturer’s instructions. Library preparation was carried out using the Quant-Seq 3’ mRNA-Seq Library Prep Kit (Lexogen, 016). Sequencing was performed on the HiSeq4000 with a SE65 run at the UCSF Center for Advanced Technologies (CAT).

#### Analysis of mRNA decay rate

We first used Salmon (v0.14.1) to align the reads to the human transcriptome (gencode v28). Transcripts with average expression >1 TPM across the cell lines were selected for further analysis (i.e. transcripts that are ‘expressed’). Bowtie2 (v2.3.5) was then used to align reads to these annotated major isoforms. Samtools mpileup command was used next to generate read stacks, which were then processed to count the number of T to C conversions. For every gene, we required the coverage to be at least 10 reads at counted Ts, filtered abundant variant calls (as they imply germline variant rather than chemical conversion), and finally calculated a C to T ratio (allowing for a pseudo-count of 1). We filtered outlier transcripts, using the 3×IQR above or below median as thresholds. Note that biological replicates were combined at this stage by combining the counts prior to calculating the C to T ratio. Since the cells were pulsed with 4sU for 4hr, 4sU incorporation, and therefore the resulting C to T ratio, are driven by the rate of transcription. At steady state, decay rate is equal to transcription rate normalized by the steady state expression. So, we used this rule to estimate a decay rate (log transformed) for every gene across all cell lines, based on its SLAM-seq C to T ratio and its baseline expression (AU).

### Modeling context-specific mRNA stability using GreyHound

#### Data preparation and pre-processing

RNA expression data from breast cancer cell lines (HCC1806, MDA-MB-231, MCF7, MDA-MB-453, HCC38, and ZR-75) from above was normalized in transcripts per million (TPM) and log-transformed. Genes with low expression (sum of expression values across all samples ≤ 10) were filtered out, resulting in 12,388 genes for downstream analysis. RNA binding proteins (RBPs) were identified using the Gene Ontology term "RNA binding" (GO:0003723), which yielded 1,378 RBPs that were present in our expression dataset. RNA decay estimates were similarly collected as described above.

For sequence data, we extracted the major isoform sequences from GENCODE v28, limiting our analysis to transcripts between 512 and 4,096 nucleotides in length. This resulted in ∼7,000 transcripts with both sequence and expression data. The dataset was split into training (80%), validation (5%), and testing (15%) sets.

#### Model architecture

We developed a hybrid deep learning model, named GreyHound, that integrates sequence information and cell-type-specific RNA binding protein (RBP) expression to predict transcript stability. The model consists of three main components:

1. **Sequence Module**: A convolutional neural network (CNN) that processes transcript sequences. The sequences were one-hot encoded across four channels (A, C, G, T) and passed through two convolutional blocks. Each block included a one-dimensional convolutional layer, leaky ReLU activation, batch normalization, max pooling, and dropout layers. Following these blocks, three residual parallel dilated convolutional layers with dilation rates of 1, 2, and 4 were applied to capture sequence features at different scales.
2. **RBP Expression Module**: A variational autoencoder (VAE) pre-trained on RBP expression data. The encoder component of the VAE compresses the 1,378-dimensional RBP expression vector into a 50-dimensional latent space, capturing the essential patterns of RBP expression across cell types.
3. **Integration Module**: A fully connected neural network that combines the flattened outputs from the sequence CNN and the latent representation from the RBP VAE to predict transcript stability.

#### VAE pre-training

First, a VAE was trained to encode the RBP expression profiles. The VAE consisted of an encoder with a 500-neuron hidden layer and a latent space of 50 dimensions. The loss function combined a reconstruction loss (mean squared error) and a KL divergence term with a gradually increasing weight (β) to promote disentanglement of the latent space. The quality of the VAE was evaluated by calculating the Pearson correlation between the original expression values and the reconstructed values.

#### GreyHound model training

The encoder and mu (mean) components of the pre-trained VAE were then transferred to the GreyHound model, with their weights initially frozen. The model was trained using mean squared error (MSE) loss and the Adam optimizer with a learning rate of 0.001. Gradient clipping was applied to prevent exploding gradients.

Training proceeded in two stages:

1. Initial training for 10 epochs with the VAE encoder weights frozen
2. Fine-tuning for an additional 5 epochs with all model parameters, including the VAE encoder, unfrozen

The model was trained using a batch size of 32. During training, the model’s performance was evaluated on the validation set after each epoch, and the model with the highest Pearson correlation coefficient on the validation set was selected as the final model.

#### Model evaluation

The performance of the model was evaluated on the held-out test set using Pearson correlation between predicted and observed RNA stability values. The model achieved a Pearson correlation coefficient of 0.624 on the test set, indicating a strong ability to predict transcript stability from sequence and RBP expression data.

#### Interpretability analysis

To understand which sequence elements and RBPs contribute to stability predictions, we performed a saliency analysis. For each transcript, we calculated the gradient of the model’s output with respect to both the input sequence and RBP expression values.

For sequence interpretation, we identified regions with high gradient variance across cell lines, indicating sequences that differentially affect stability depending on cellular context. These salient regions were extracted and analyzed using FIRE (Finding Informative Regulatory Elements) to identify enriched motifs.

For RBP interpretation, we identified RBPs with high gradient values, suggesting their importance in determining variation in transcript stability. We then constructed a network connecting enriched motifs to RBPs that are likely to bind them, based on the correlation between RBP expression and the stability of transcripts containing specific motifs.

This integrative approach allowed us to identify key sequence motifs and RBPs that modulate RNA stability in a cell-state-specific manner, providing insights into post-transcriptional regulatory mechanisms in breast cancer cell lines.

### Clinical Association Studies

For survival analyses, we investigated the relationship between the expression of select RBPs (identified as important in our model) and patient outcomes in TCGA or METABRIC breast cancer datasets. Three types of survival endpoints were analyzed: disease-free survival (DFS), overall survival (OS), and progression-free survival (PFS).

For each survival analysis, we applied the following methodology:

1. Patients with missing survival data were excluded from the analysis.
2. Survival times were capped at a maximum follow-up period (120 months for DFS and OS, 60 months for PFS) to minimize the effect of outliers and focusing on clinically relevant timeframes.
3. For each RBP of interest, we systematically tested a range of expression thresholds to stratify patients into "high" and "low" expression groups. The optimal threshold was determined by selecting the cutpoint that yielded the most statistically significant difference in survival curves between groups, as assessed by the log-rank test.
4. Kaplan-Meier curves were generated for the resulting patient groups, and differences were quantified using the log-rank test. Hazard ratios (HR) were calculated as the ratio of observed to expected events in the high-expression group divided by the same ratio in the low-expression group.
5. To adjust for potential confounding variables, Cox proportional hazards models were fitted with RBP expression status as the primary variable of interest, along with clinical covariates including tumor subtype, pathological stage, age, and prior diagnosis history.

Additionally, we evaluated associations between RBP expression and clinicopathological features, such as tumor stage, using analysis of variance (ANOVA) followed by Tukey’s honestly significant difference (HSD) test for multiple comparisons.

### Cell Culture

Human breast cancer cell lines MDA-MB-231 (MDA-parental, ATCC HTB-26), MDA-LM2 (MDA-MB-231 highly metastatic derivative)^25^, MDA-MB-453 (ATCC HTB-131), MCF7 (ATCC HTB-22), HEK293T (293T, ATCC CRL-3216) and MDA-RBMS3-FLAG-tagged cells were all cultured in DMEM supplemented with 10% fetal bovine serum (FBS), penicillin (100U/mL), streptomycin (100µg/mL), and amphotericin B (1µg/mL). The HCC1806 (ATCC CRL-2335), HCC38 (ATCC CRL-2314), and ZR-75-1 (ATCC CRL-1500) human breast cancer cell lines were cultured in RPMI-1640 medium supplemented with 10% FBS, penicillin (100U/mL), streptomycin (100µg/mL), and amphotericin B (1µg/mL). Cell lines were routinely screened for *Mycoplasma* contamination with a PCR-based assay. All cell lines were grown in a humidified incubator at 37°C with 5% CO_2_.

#### Knockdown Cell Lines

The shRNAs shRBMS3-1 (TRCN0000152525) and shRBMS3-2 (TRCN0000152012) were cloned into plasmid pLKO.1-Puro (Addgene, #8453) with the plasmid pLKO.1-Puro-Scramble (Addgene, #162011) acting as negative control. These plasmids were then cotransfected with lentiviral packaging plasmids pCMV-R8.91 (Addgene, #202687) and pMD2.G (Addgene, #12259) into HEK293T cells using the TransIT-Lenti Transfection Reagent (MirusBio, MIR 6600) according to manufacturer’s instructions for lentivirus production. Virus was collected and filtered through a 0.45µm filter and 2mL viral media mixed with polybrene at final concentration 8µg/mL (Sigma-Aldrich, TR-1003) was applied to transduce luciferase-labelled MDA-MB-231 and HCC1806 cell lines. Virus was removed after 8hrs and cells were rested for 72hrs. For selection, puromycin was applied at 1µg/mL until control cells (no virus) were all dead. Knockdown was assessed via qPCR.

For dual shRNA constructs, gene blocks of the dual shRNA cassettes containing either shScramble1-shScramble2, shScramble1-shTXNIP, or shRBMS3-1-shTXNIP were ordered from TwistBio. The shRNAs used to generate these gene blocks were the following: shTXNIP (TRCN0000262802), shRBMS3-1 (TRCN0000152525), and 2 scramble sequences. These gene blocks were then cloned into the pLKO.1-Puro (Addgene, #8453) plasmid and then transduced as described above into luciferase labelled MDA-MB-231 cells.

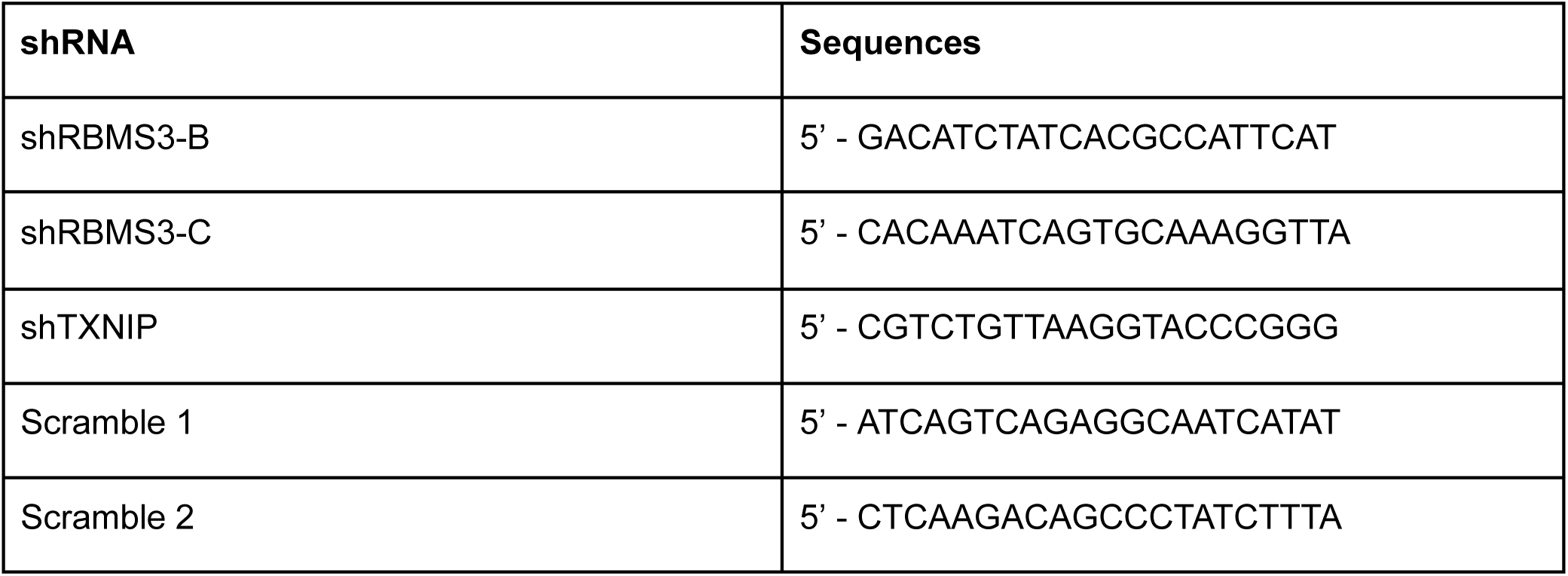

#### Overexpression Cell Lines

RBMS3 overexpression cells were generated by stably transducing the pLX-304 lentiviral vector (Addgene, #25890) containing the RBMS3 open reading frame into MDA-LM2 cells.

#### RBMS3 Flag-Tagged MDA-MB-231

RBMS3 Flag-tagged cells were generated by stably transducing the lentiviral vector pLX302EF1a containing RBMS3 + N-ter 2xFLAG fusions into MDA-MB-231 breast cancer cells.

### RNA isolation and RT-qPCR

Total RNA was isolated from cells using TRIzol Reagent (Invitrogen, 15596018) according to the manufacturer’s instructions. cDNA synthesis took place with Maxima H Minus Reverse Transcriptase (Thermo Scientific, EP0751) and for qPCR, PerfeCTa SYBR Green SuperMix (QuantaBio, 95056-500) was used with HPRT1 as endogenous control.

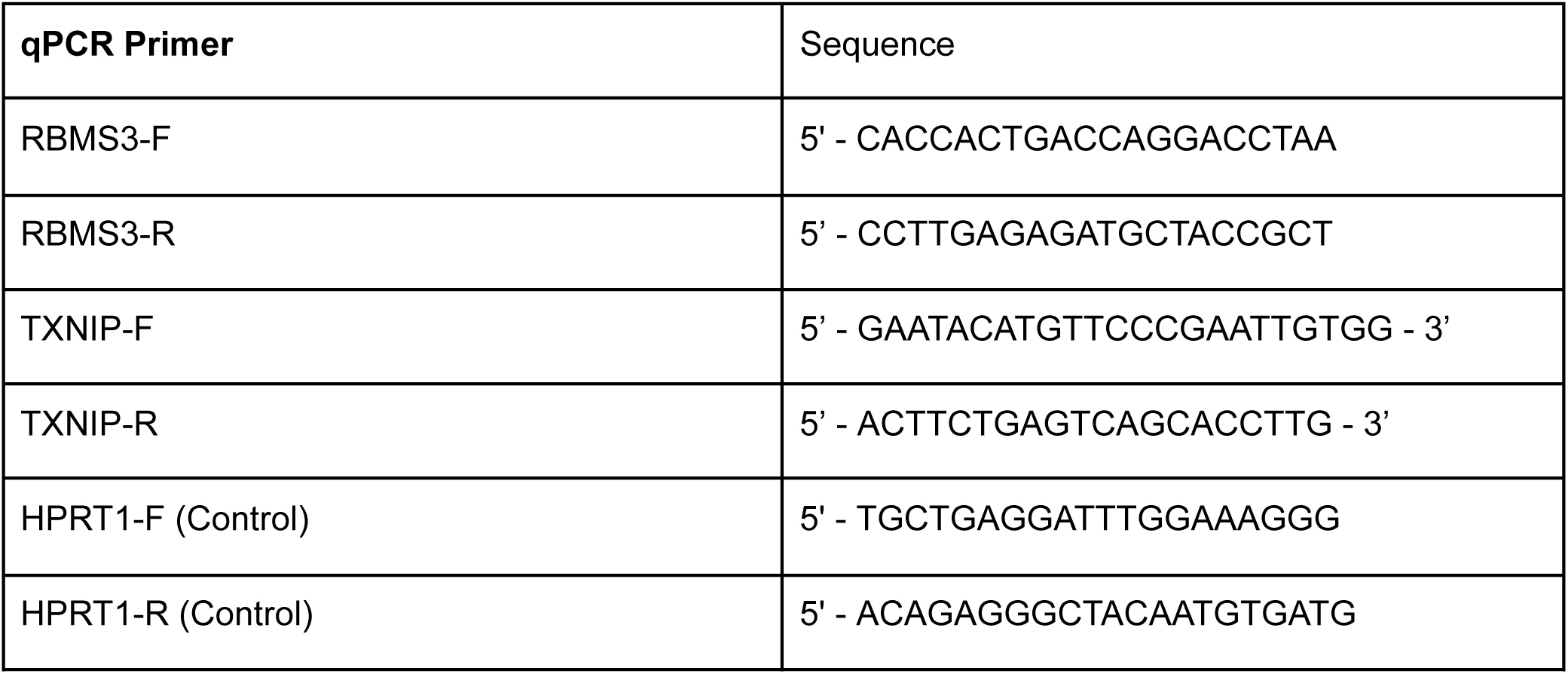

### Total RNA-sequencing

RNA was isolated from MDA-MB-231 cells with RBMS3 knockdown and control in biological replicates using TRIzol reagent as described above. Libraries for RNA-seq were prepared using the SMARTer Stranded Total RNA-Seq Kit v3 - Pico Input Mammalian (Takara, 634485). Purified libraries were quantified on the TapeStation with D1000 HS tape (Agilent, 5067-5585). Sequencing was performed on Illumina NextSeq 500 with a PE75 run in Helen Diller Cancer Center.

### RBMS3 CLIP-seq

#### Cell Crosslinking

For crosslinking, MDA-MB-231 cells containing FLAG-tagged RBMS3 were harvested at ∼90% confluency from 4-15cm plates per replicate. On ice, cells were rinsed with 1X PBS, then in 1X PBS, cells were irradiated using a 254nm crosslinker set to 400mJ/cm2 UV. Plates were placed back on ice and cells scraped, collected into tubes, spun down at 2000xg at 4°C for 2 minutes, and then the remaining PBS was removed. Pellets were then frozen at -80°C.

#### Immunoprecipitation

Cell pellets were resuspended and lysed on ice for 10 minutes in 300µL per plate of FLAG Lysis Buffer (25mM Tris pH7.5, 150mM NaCl, 1% IGEPAL CA-630, and 5% Glycerol) plus 3µL of 100X Protease inhibitor and 3µL of RNAse inhibitor. DNaseI was added at 30µL per plate to lysate and incubated at 37°C for 10 minutes shaking at 1000rpm. Lysate was then divided equally into 2 tubes. One tube is for 10µL per plate of High RNase mix (RNaseA 1:3,000 + RNaseI 1:100) and the other tube is for 10µL per plate of Low RNase mix (RNaseA 1:15,000 + RNaseI 1:500). Lysate is then incubated at 37°C for 5 minutes. The two tubes are then combined and spun down at 4°C at ∼20,000xg for 20 minutes. FLAG Magnetic Agarose Beads (Pierce Anti-DYKDDDDK #A36797) that were pre-washed twice in PBST (0.02% tween-20) and once in Low Salt Wash Buffer (1X PBS with no Mg++ and no Ca++, 0.1% SDS, 0.5% Sodium Deoxycholate, 0.5% IGEPAL CA-630) were then used for immunoprecipitation at 12.5µL of beads per plate. Cleared supernatant wasthen added to beads and rotated end-over-end for 1-2 hours at 4°C. Beads were then washed twice with cold low salt wash buffer, twice with cold high salt wash buffer (5X PBS with no Mg++ and no Ca++, 0.1% SDS, 0.5% Sodium Deoxycholate, 0.5% IGEPAL CA-630), and twice with cold PNK wash buffer (50mM Tris-HCl pH 7.5, 10mM MgCl2, 0.5% IGEPAL CA-630).

#### Dephosphorylation

Immunoprecipitated RNA-Protein (RNP) complexes were dephosphorylated on-beads by resuspending beads in 2.5µL 10X PNK buffer (500mM Tris pH 6.8, 5mM MgCl2, 50mM DTT), 2µL T4 PNK (10u/µL, NEB), 0.5µL RNase inhibitor, 20µL nuclease free water. Samples were then incubated at 37°C for 20 minutes, shaking at 1350rpm 15 seconds /5 minutes rest. Beads were then washed once with cold PNK wash buffer, once with cold high salt wash buffer (>1 minute) and twice with PNK wash buffer.

#### PolyA tailing RNP complexes

RNP complexes undergo polyA-tailing following dephosphorylation. On ice beads were resuspended in 0.8µL of Yeast PAP (Jena 600U/µL), 4µL 5X yeast PAP buffer, 10mM ATP (unlabeled), 0.5µL RNase inhibitor, and 13.7µL nuclease free water. Samples were then incubated at 22°C for 5 minutes and with one 15 second shake at 1350rpm. Beads were then washed twice with cold high salt wash buffer and twice with cold PNK wash buffer.

#### N3-dUTP end labeling of RNP complexes

To N3-dUTP end labeled RNA, the beads were resuspended in 0.4µL Yeast PAP (Jena 600U/µL), 2µL 5X yeast PAP buffer, 0.25µL RNase inhibitor, 2µL 10mM N3-dUTP, and 5.35µL nuclease free water. Samples were then incubated at 37°C for 20 minutes with shaking at 1350rpm 15 seconds / 5 minutes rest. Beads were then washed twice in cold high salt wash buffer and twice in cold 1X PBS

#### Dye labeling of N3-labeled RNP complexes

RNPs were then labeled by resuspending beads in 20µL 800CW DBCO dye (0.2mM in 1XPBS). Beads were incubated at 22°C for 30 minutes with shaking at 1350rpm for 15 seconds / 5 minutes rest. Beads were then washed once with cold high salt wash buffer and once with cold PNK wash buffer. Then beads were resuspended in loading buffer (1X NuPAGE loading buffer, 50mM DTT, diluted in PNK wash buffer) and incubated at75°C for 10 minutes shaking at 1000rpm protected from light. Samples were then placed on magnet and eluate removed to clean tubes.

#### PAGE Gel and Transfer

Eluted samples were run on a 12-well Novex NuPAGE 4-12% Bis-Tris gel (1mm thick) at 180V in 1X MOPS running buffer at 4°C, protected from light. Samples were transferred to a Protran BA-85 nitrocellulose membrane in 1X NuPAGE transfer buffer with 10% ethanol at 30V for 75 minutes. Membrane was imaged on a LI-COR Odyssey imaging instrument. The region of the membrane containing the RNA-protein complexes was then cut out and placed in a clean tube.

#### Proteinase K digest and RNA capture

RNA was isolated from the membrane by digesting protein in 200µL protein K digestion buffer (100mM Tris-HCL pH7.5, 100mM NaCl, 1mM EDTA, 0.2% SDS), and 12.5µL proteinase K at 55°C for 45 minutes shaking at 1100rpm. To capture RNA, the supernatant was then transferred to a clean tube and adjusted to 0.5M NaCl. This solution was then added to oligo d(T) magnetic beads pre-washed twice with proteinase K buffer. Samples were then incubated at 25°C for 20 minutes shaking at 300rpm with 1350rpm increases for 10 seconds every 10 minutes. Beads were then washed twice with cold high salt wash buffer and twice with cold 1X PBS. Samples are then eluted off of the beads in 8µL of TE elution buffer (20mM Tris-HCl pH 7.5, 1mM EDTA) and at 50°C for 5 minutes. Supernatant was transferred to PCR tubes in preparation for cDNA synthesis.

#### cDNA synthesis

For CLIP-seq library synthesis, the Takara SMARTer smRNA-Seq Kit was used with some modifications to the manufacturer’s protocol. Briefly, to the captured RNA from the previous step, 2.5µL smRNA Mix 1 and 10uM UMI RT primer was added and incubated at 72°C for 3 minutes followed by 2-5 minutes on ice. Then 6.5µL smRNA Mix 2, 0.5µL RNase inhibitor, and 2µL PrimeScript RT (200U/µL) was added and samples incubated at 42°C for 60 minutes, 70°C for 10 minutes, and then placed back on ice.

#### PCR amplification

To the 20µL of cDNA from the previous step, 50µL of 2X SeqAmp CB PCR Buffer, 2µL SeqAmp DNA Polymerase, 2µL 10uM Universal Reverse Primer and 24µL nuclease free water was added. Then 2µL of a unique 10µM indexed TruSeq forward primer is added separately to each sample. Samples were amplified using the following thermocycler program: 98°C 1 minute for 1 cycle, 98°C 10 seconds - 60°C 5 seconds - 68°C 10 seconds for 18 cycles, and hold at 4°C. Libraries were size selected using Zymo Select-a-Size MagBeads at 1.1X bead to PCR ratio according to manufacturer’s instructions. Library size distribution and quantity was assessed using the Agilent D1000 ScreenTape System and an Agilent Tapestation 4200 according to manufacturer’s instructions. Sequencing was performed on a HiSeq 4000 at the UCSF Center for Advanced Technologies.

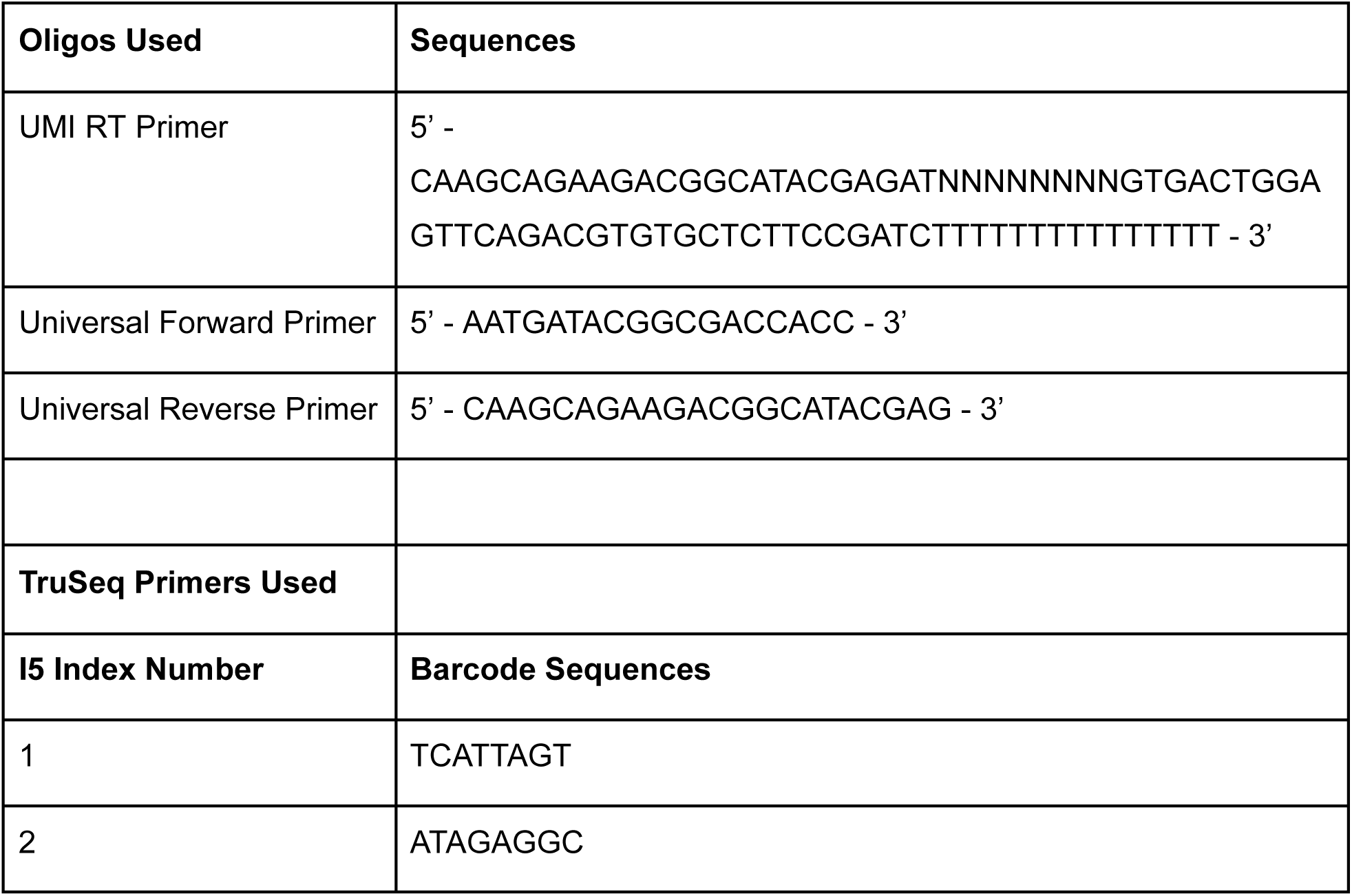

#### CLIP-seq data analysis

Raw sequencing data from RBMS3-1 and RBMS3-2 samples were first processed by appending unique molecular identifiers (UMIs) to reads using a custom Python script (append_umi.py).

The processed reads were then analyzed using the CLIP Tool Kit (CTK) pipeline. The CTK pipeline included alignment to the human reference genome (hg38), filtering, and peak calling. CIMS (Crosslinking-Induced Mutation Sites) analysis was performed to identify nucleotide deletions that represent direct protein-RNA crosslinking sites. These analyses resulted in 27,706 high-confidence RBMS3 binding sites (FDR < 10%, fold change > 10) that were stored in the BED format file RBMS3_eclairCLIP.pool.tag.uniq.del.CIMS.fdr10.f10.bed.

To characterize the distribution of RBMS3 binding sites across genomic features, we performed systematic intersection of the binding sites with genomic annotations using BEDTools. The binding sites were categorized into 3’ UTRs, 5’ UTRs, coding sequences (CDS), introns, and intergenic regions by sequential subtraction:

1. 3’ UTR peaks were directly intersected with the hg38 3’ UTR annotation.
2. 5’ UTR peaks were identified by excluding 3’ UTR peaks and intersecting with 5’ UTR annotations.
3. CDS peaks were identified by excluding 3’ and 5’ UTR peaks and intersecting with exon annotations.
4. Intronic peaks were identified by excluding all previous categories and intersecting with intron annotations.
5. Intergenic peaks were calculated as the remaining sites.

We found that RBMS3 binding was predominantly enriched in 3’ UTRs (73.3% of peaks), with smaller fractions in introns (10.9%), intergenic regions (12.1%), CDS (2.5%), and 5’ UTRs (1.1%). This distribution was visualized using the UpSetR package in R, which illustrated the overlaps between binding sites in various genomic features.

To identify sequence motifs recognized by RBMS3, we extracted the sequences corresponding to the binding sites using twoBitToFa with the hg38 reference genome. The sequences were analyzed using FIRE (Finding Informative Regulatory Elements), a mutual information-based motif discovery algorithm. Two approaches were employed:

1. Non-discovery mode: Using a previously defined RBMS3 motif pattern [AGT][AT]ATA[AT]A to validate its enrichment in our dataset.
2. Discovery mode: De novo identification of enriched sequence patterns.

FIRE analysis confirmed strong enrichment of A/U-rich motifs, with the most significant being the canonical AATAAA polyadenylation signal (z-score = 1939.19). Additional enriched motifs included several variants of A/T-rich sequences, suggesting RBMS3’s preference for binding to such regions.

### Protein label-free quantification

In preparation for whole protein analysis, RBMS3 knockdown and control MDA-MB-231 cells were washed 3x with PBS, scraped, pelleted, and snap frozen. After thawing stored pellets on ice, samples were resuspended in lysis buffer (PreOmics) supplemented with 20mg/mL complete mini protease inhibitor cocktail (Sigma Aldrich) before preparation with iST96 kit (PreOmics). Cell lines were collected in biological triplicates and samples were run in technical duplicates on the mass spectrometer.

A nanoElute system was attached in line to a timsTOF Pro mass spectrometer equipped with a CaptiveSpray source (Bruker). Chromatography was conducted at 40°C through a 25cm reversed-phase C18 PepSep column (Bruker) at a constant flow rate of 0.5 μL/min. Mobile phase A was 98/2/0.1% Water/MeCN/Formic Acid (v/v/v) and phase B was MeCN with 0.1% Formic Acid (v/v). During a 108 min method, peptides were separated by a 3-step linear gradient (5% to 30% B over 90 min, 30% to 35% B over 10 min, 35% to 95% B over 4 min) followed by a 4 min isocratic flush at 95% for 4 min before washing and a return to low organic conditions. Experiments were run as data-dependent acquisitions with ion mobility activated in PASEF mode. MS and MS/MS spectra were collected with m/z 100 to 1700 and ions with z = +1 were excluded.

Raw data files were searched using PEAKS Online Xpro 1.6 (Bioinformatics Solutions Inc.). The precursor mass error tolerance and fragment mass error tolerance were set to 20 ppm and 0.03 respectively. The trypsin digest mode was set to semi-specific and missed cleavages was set to 2. The human Swiss-Prot reviewed (canonical) database (downloaded from UniProt) and the common repository of adventitious proteins (cRAP, downloaded from The Global Proteome Machine Organization) totaling 20,487 entries were used. Carbamidomethylation was selected as a fixed modification. Oxidation (M) was selected as a variable modification.

### Parallel reporter assays

#### Generation of dual reporter library

We performed a reporter assay by introducing the targets’ respective RBMS3 binding site sequences into the dual-promoter pBdLV-Puro-T2A-mCherry vector previously described by Yu et al. (2020, supplementary file 8)^10^. First, we selected RBMS3 binding site sequences of the listed targets based on the magnitude of ecCLIP peaks within the 3’UTR of the transcripts. For each binding site, we generated paired scrambled control sequences that preserve the dinucleotide composition using *ushuffle*^32^. Then, DNA oligonucleotides (116 bp long) were synthesized (IDT DNA) for these paired sequences and cloned into the pBdLV-Puro-T2A-mCherry reporter vector^10^ through PacI digestion (NEB, R0547S) and HiFi assembly (NEB, E2621S) in an arrayed format. The plasmid library was pooled in an equimolar ratio and lentivirus was prepared using TransIT-Lenti (MirusBio, MIR 6600) in HEK293T cells. Next, we transduced the viral library into MDA-MB-231 RBMS3 knockdown and control cells for 8h, rested the cells and sorted after 72h for mCherry positive cells. For library preparation, we used the DNA/RNA co-extraction (Zymo Research, D7001), followed by reverse transcription of RNA samples with the following primer: 5’ CTCTTTCCCTACACGACGCTCTTCCGATCTNNNNNNNNNNNtggtctggatccaccggtccgg targeting GFP. Targeted amplification of gDNA and cDNA for mCherry (internal control for transcription rate) and GFP (indicative of mRNA stability) including the binding site sequences was performed with the primers listed below. Sequencing of amplified DNA and cDNA libraries was performed on HiSeq 4000 at the UCSF Center of Advanced Technologies (CAT).

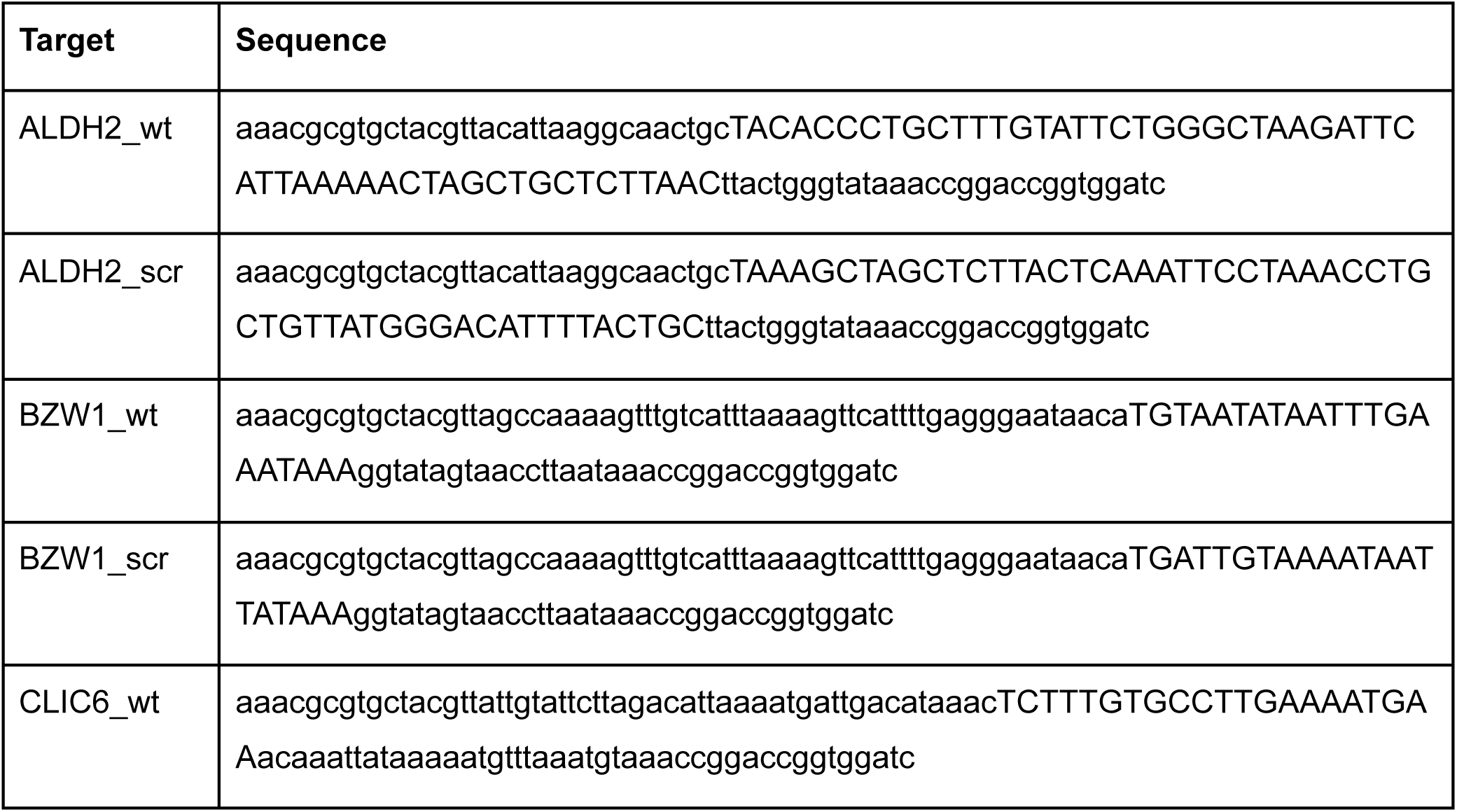
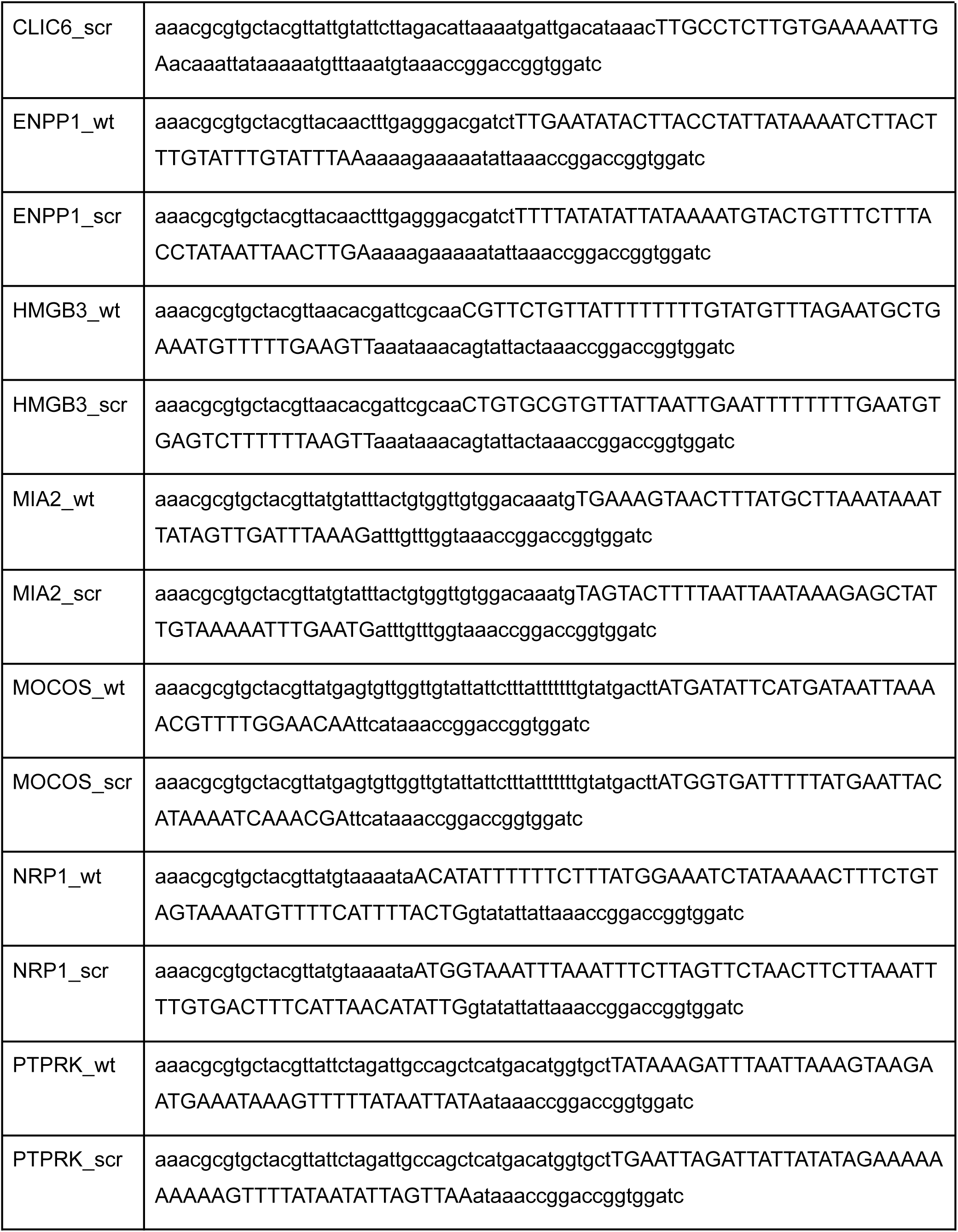
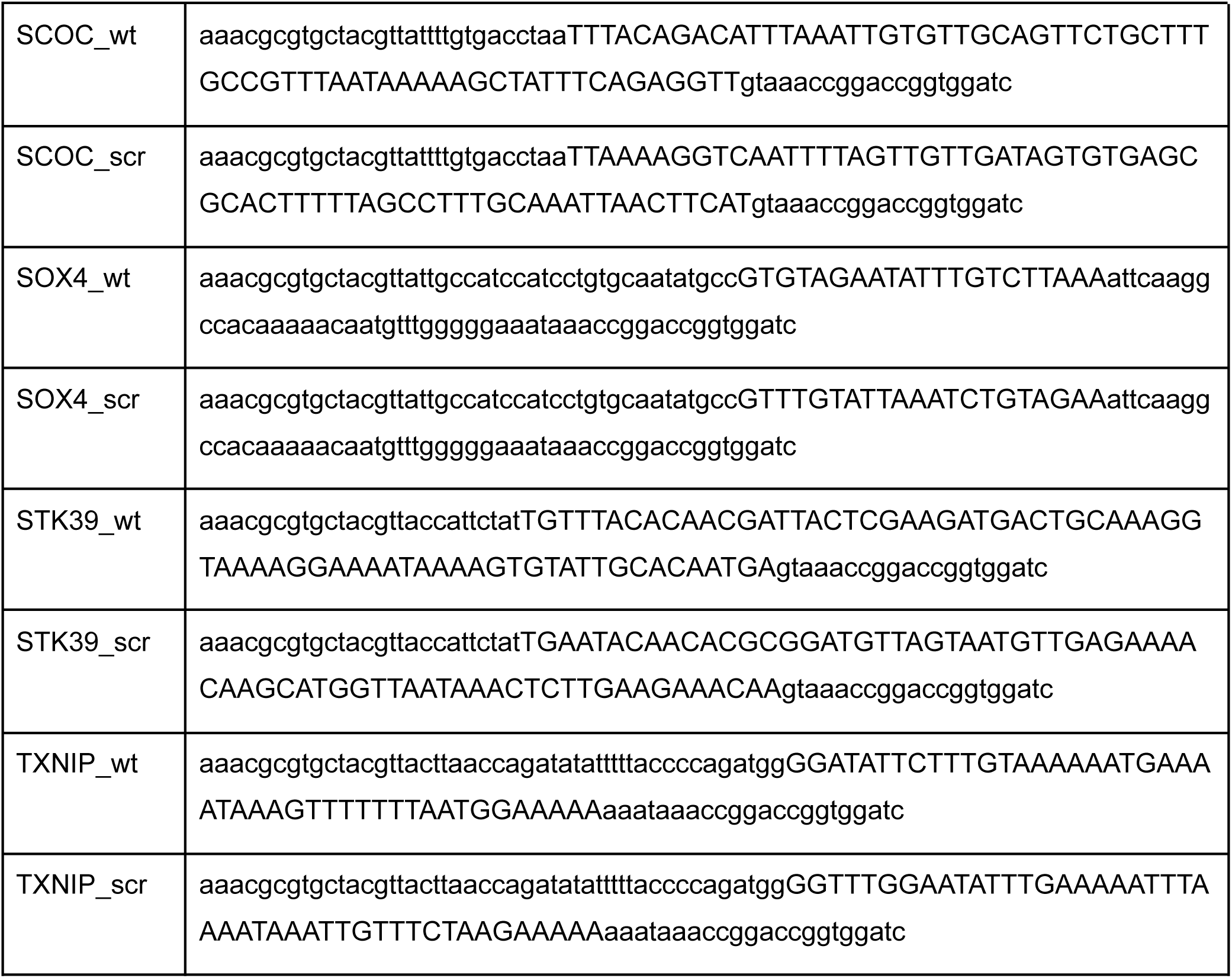

#### Analysis of the parallel reporter assay data

We first collated sequence reads with identical sequences using the fastq2collapse.pl program (CTK tools) to remove PCR duplicates, generating collapsed FASTQ files. UMIs were extracted from these collapsed sequences using UMI-tools. The adapter sequence (TGGTCTGGATCCACCGGTCCGGT) was then removed from the UMI-extracted reads using cutadapt (parameters: --trimmed-only -e 0.2 -g TGGTCTGGATCCACCGGTCCGGT -m 25), retaining only reads that contained the adapter and were at least 25 nucleotides long after trimming.

The processed RNA reads were then reverse-complemented using seqkit to align them with the reference sequences. For DNA libraries, a similar processing pipeline was used, but with a different adapter sequence (GTGGTCTGGATCCACCGGTCCGGTTTA), and a minimum length filter of 35 nucleotides.

A reference sequence file containing all candidate regulatory elements was indexed using BWA. Both RNA and DNA processed reads were aligned to this reference using BWA-MEM. Aligned reads were sorted and indexed using SAMtools. For RNA libraries, we performed UMI-based deduplication to account for PCR duplicates by collapsing reads with identical UMIs mapping to the same position using UMICollapse (parameters: -k 1 --algo adj --merge avgqual). Counts for each candidate regulatory element were obtained by extracting the reference sequence names from the alignment files and counting their occurrences using SAMtools and standard Unix tools.

We used DESeq2 for differential expression analysis. Size factors were calculated from the scrambled (SCR) sequences to normalize for sequencing depth differences. We performed two main comparisons:

1. For shCTRL samples, we tested the expression difference between reference (REF) and scrambled (SCR) sequences by calculating log2 fold changes and adjusted p-values.
2. For shRBMS3 samples, we performed the same comparison between reference and scrambled sequences.

The design formula for DESeq2 represented the combination of sequence type (REF/SCR) and library type (RNA/DNA). We used contrast lists to specifically test the interaction between sequence type and library type, effectively measuring the regulatory activity of the sequences while controlling for differences in DNA abundance. Statistical significance of the overall difference in regulatory activity between control and RBMS3 knockdown conditions was assessed using a Wilcoxon rank-sum test.

### Animal Studies

All animal studies were performed according to IACUC guidelines (IACUC approval number AN194337-01L). Metastatic lung colonization assays were done using age-matched female NOD scid gamma mice (Jackson Labs, 005557). *In vivo* bioluminescence was used to track metastasis and measured by injection of luciferin (Perkin-Elmer) followed by imaging on an IVIS instrument. Hematoxylin and eosin (H&E) staining of lung tissue sections was used for histology.

### Dual Guide CRIPSRi Assays

#### Generation of dual guide library

The dual-guide CRISPRi-screen sgRNA library, containing 13 targets and 10 non-targeting controls, was designed according to a protocol previously described and oligos were synthesized (IDT). Briefly, two sgRNAs per target gene were inserted into the dual-guide plasmid pJR85 (Addgene, 140095) as the backbone with Golden-Gate assembly from pJR89 (Addgene, 140096)^27^ using two-step cloning first with BstXI (NEB, R0113S) and BlpI (NEB, R0585S) and second with BsmBI-v2 (NEB, R0739S). The sgRNA sequences were selected from Horlbeck et al. (2016, supplementary file 3)^33^.

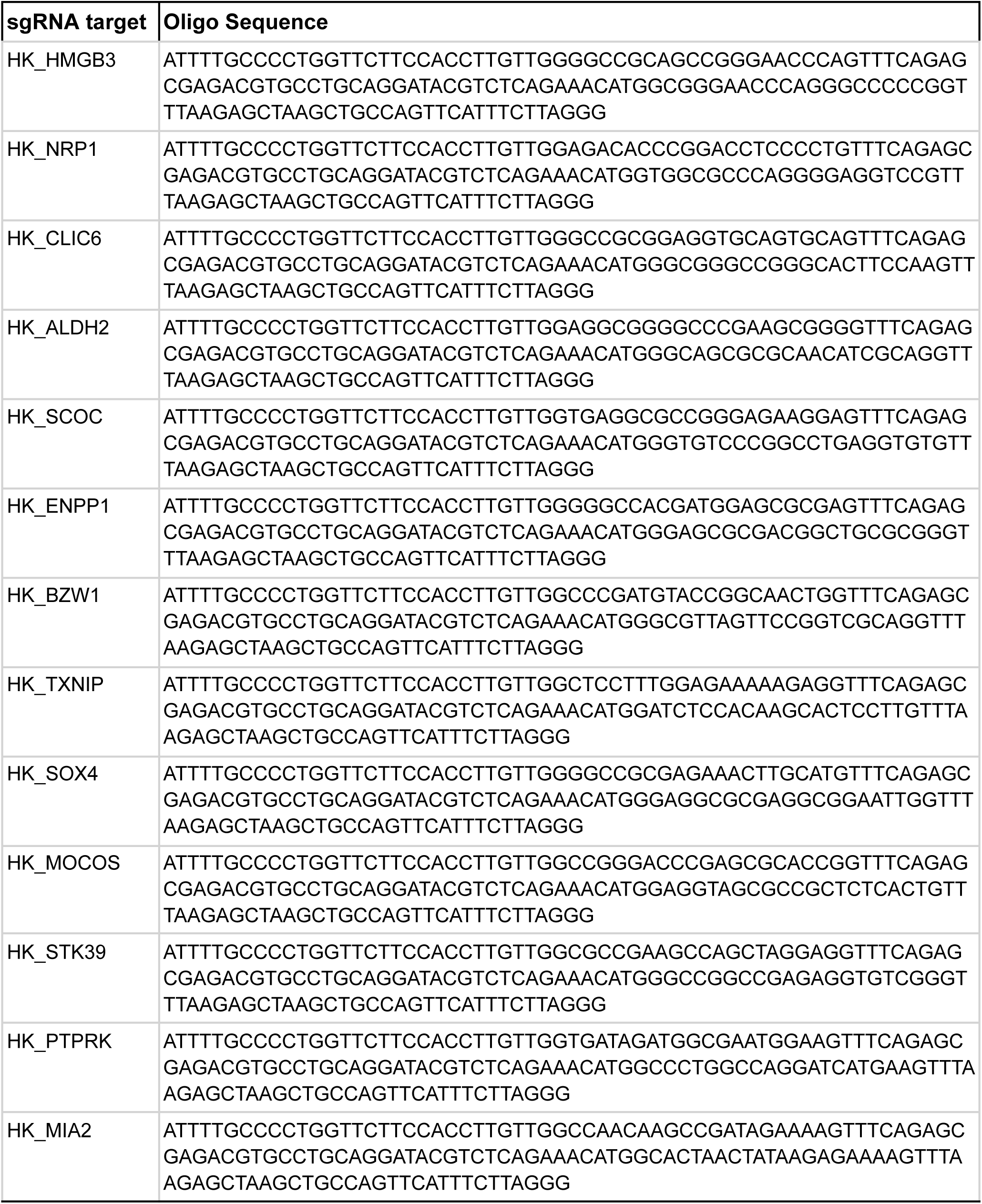
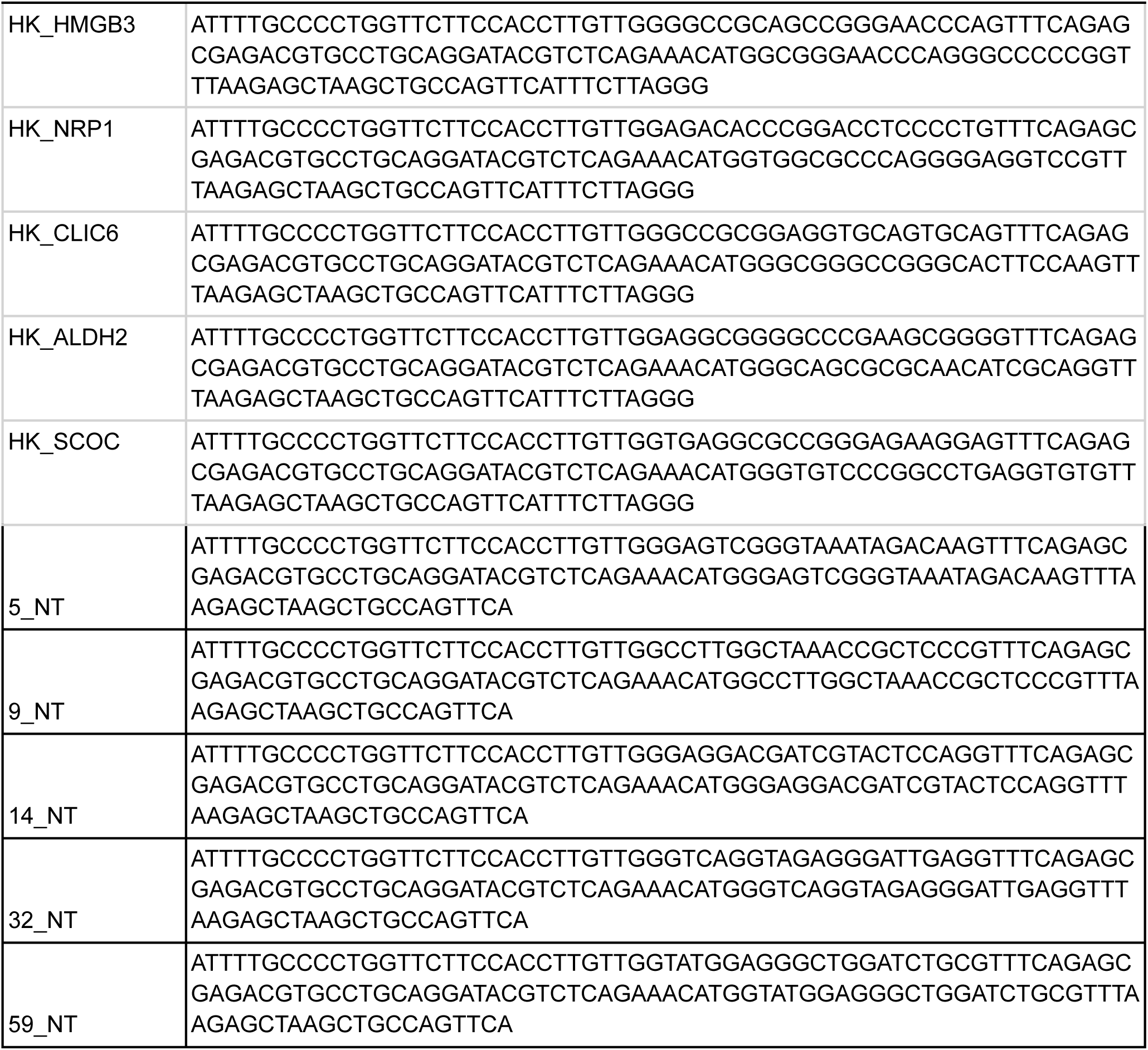

The two-step cloning was performed using an arrayed format and the resulting plasmids were pooled at an equimolar ratio. The dual sgRNA plasmid library was then transduced into Zim3-CRISPRi^34^ ready MDA-MB-231 cells via lentiviral transduction as described above. Cells were then plated (*in vitro* arm) and injected into tail veins of NGS mice (*in vivo* arm). For the *in vitro* arm, 1×10^5^ cells were plated and cells from day 0 as well as cells after 10 doublings had occurred were collected for library preparation. For the *in vivo* arm, 2.5×10^5^ cells were injected via tail vein into NGS mice and lungs were collected at ∼5 weeks. The gDNA from both experimental arms was isolated and library preparation was done according to a protocol previously described by Replogle *et al*. (2020)^27^. Sequencing was performed on NovaSeq X 1.5B with a PE100 run.

#### Analysis of the CRISPRi assay data

Sequencing data from in vitro and in vivo CRISPR screens was processed using a multi-step computational workflow. Raw FASTQ files were first trimmed to remove adapter sequences using Cutadapt (v4.9). For each sample, forward reads were trimmed for the 5’ adapter sequence (GTTTCAGAGCTAAGCACAAGAGTGCATAGCAA) and reverse reads for the 3’ adapter sequence (CCATGTTTCTGGCTTTCCACAAGATATATAAA). The trimming parameters were set to retain only reads with a fixed length of 19 nucleotides to ensure precise sgRNA sequence capture. Paired-end reads were processed together. The trimmed reads were aligned to a custom reference containing dual guide sequences using Bowtie2 (v2.3.4.2). The alignment was performed with sensitive parameters and end-to-end mode (--sensitive --end-to-end -N 1) to ensure accurate mapping of sgRNA sequences. Only properly paired reads that aligned to the reference were retained using SAMtools filtering parameters (-f 0x1 -f 0x2 -F 0x4 -F 0x8), generating BAM files for each sample.

Read counts for each sgRNA were extracted from the aligned BAM files using SAMtools. For each sample, the third field (RNAME) in the SAM format was extracted to identify the target sgRNA, and the occurrences were counted using the Unix commands sort and uniq -c. The resulting count tables were stored in tab-delimited format. The count data was processed in R, where individual count files were merged into a combined count matrix. To account for paired-end reads, the raw counts were divided by 2 to avoid double-counting. The matrix was structured with sgRNAs as rows and samples as columns, with 9 total samples representing:

- Three Day 0 replicates (RBMS3-TL-D0-1, RBMS3-TL-D0-2, RBMS3-TL-D0-3)
- Three Day 10 in vitro samples (RBMS3-TL-D10-1, RBMS3-TL-D10-2, RBMS3-TL-D10-3)
- Three in vivo tumor samples (RBMS3-TL-M-1, RBMS3-TL-M-2, RBMS3-TL-M-3)

Differential abundance analysis was performed using the DESeq2 package (v1.42.0) in R to identify sgRNAs with significant changes in abundance between conditions. Non-targeting control sgRNAs (prefixed with "NT_") were used for normalization and as a reference for calculating size factors to account for sequencing depth differences between samples.

The experimental design included three conditions (D0, D10, TL-M) with three biological replicates each. Three key comparisons were analyzed:

1. In vivo (TL-M) vs Day 0 (D0)
2. In vitro (D10) vs Day 0 (D0)
3. In vivo (TL-M) vs In vitro (D10)

Samples were grouped by condition, with Day 0 samples serving as the reference for both in vitro and in vivo comparisons. The DESeq2 analysis pipeline included estimating size factors, calculating dispersions, and fitting negative binomial models to determine log2 fold changes and adjusted p-values for each sgRNA across conditions.

A scatter plot was generated to compare the log2 fold changes of sgRNAs in the in vivo screen (x-axis) versus the in vitro screen (y-axis), both relative to Day 0. Each point represents an individual sgRNA, with color indicating the statistical significance (adjusted p-value). Significantly depleted or enriched sgRNAs (adjusted p-value < 0.01) were labeled on the plot. This visualization allowed for direct comparison of sgRNA behavior in the two screening conditions and identification of context-specific essential genes.

### Proliferation Assays

On day 0, 5×10^4^ cells per well were seeded onto 6-well plates in biological triplicates. Cells were then trypsinized, collected, and stained with 0.4% Trypan Blue Solution (Gibco) to determine cell viability on day 3 and day 5. The number of viable cells was counted using a TC20 Automated Cell Counter (Bio-Rad). A linearized exponential growth model (*ln*(*N*_*t*_) = *ln*(*N*_0_) + *r* × *t*, with time *t* [days], proliferation rate *r* [day^-1^] and number of cells *N*) was used to fit a proliferation rate for each cell line. To compare proliferation rates and test for significant differences, an unpaired two-sided t-test was used.

### Invasion Assays

For invasion assays, RBMS3 knockdown and control MDA-MB-231 cells were seeded into Corning BioCoat Matrigel Invasion Chambers with 8.0μm PET membranes (Corning) at a final concentration of 8×10⁴ cells per chamber in 500 μl of serum-free medium. Then, 750μL of complete culture medium containing 10% FBS, as a chemoattractant, was added to the lower chamber. The cells were incubated at 37°C for 24 hours. After incubation, the inserts were gently washed three times with cold PBS, and the cells were fixed using 100% methanol. Non-migrated cells on the top of the inserts were removed using cotton swabs. Following fixation, the cells that had migrated to the bottom of the inserts were washed again with PBS and stained with 0.4% crystal violet. Migrated cells were counted from three random fields per insert at a magnification of 20×, and the results were averaged from at least three biological replicates.

### Measurements in Clinical Samples

We measured TXNIP expression using qPCR in 96 clinical samples across all stages of breast cancer, namely 5 normal epithelial, 23 stage I, 30 stage II, 29 stage III, and 9 stage IV metastatic biopsies (Origene, BCRT102, BCRT103), from which 90 samples yielded sufficient amount of cDNA. HPRT was used as an endogenous control and relative TXNIP expression levels were measured using the primers listed below.

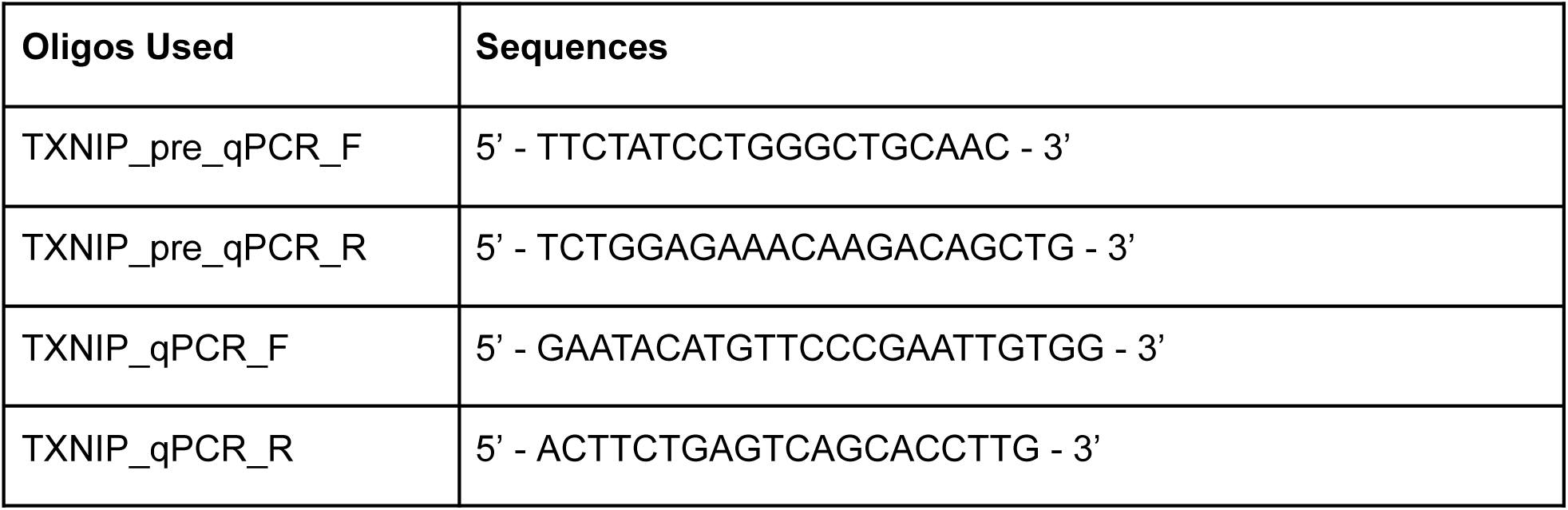

